# Nucleotide limitation results in impaired photosynthesis, reduced growth and seed yield together with massively altered gene expression

**DOI:** 10.1101/2021.01.22.427776

**Authors:** Leo Bellin, Michael Melzer, Alexander Hilo, Diana Laura Garza Amaya, Isabel Keller, Jörg Meurer, Torsten Möhlmann

**Author notes:** **Author for contact:** Dr. Torsten Möhlmann, Universität Kaiserslautern, Pflanzenphysiologie, Postfach 3049, D-67653 Kaiserslautern, Germany. **AUTHOR CONTRIBUTIONS:** T.M. conceived and supervised the study, obtained funding, and provided resources. L.B. generated all mutants and performed characterization, M.M. performed morphological and ultrastructure analysis, A.H. performed metabolite measurements, D.L.G.A. performed growth, carbohydrate and gene expression analysis, I.K. advised and interpreted ROS determination, J.M. advised and interpreted determination of photosynthesis parameters. T.M. and L.B. wrote the original draft. All authors reviewed and edited the manuscript.

## Abstract

Nucleotide limitation and imbalance is a well described phenomenon in animal research but understudied in the plant field. A peculiarity of pyrimidine de novo synthesis in plants is the complex subcellular organization. Here, we studied two organellar localized enzymes in the pathway, with chloroplast aspartate transcarbamoylase (ATC), and mitochondrial dihydroorotate dehydrogenase (DHODH). *ATC* knockdowns were most severely affected, exhibiting low levels of pyrimidine nucleotides, a low energy state, reduced photosynthetic capacity and accumulation of reactive oxygen species (ROS). Furthermore, altered leaf morphology and chloroplast ultrastructure were observed in *ATC* mutants. Although less affected, *DHODH* knockdown mutants showed impaired seed germination and altered mitochondrial ultrastructure. Transcriptome analysis of an *ATC*-amiRNA line revealed massive alterations in gene expression with central metabolic pathways being downregulated and stress response and RNA related pathways being upregulated. In addition, genes involved in central carbon metabolism, intracellular transport and respiration were mainly downregulated in ATC mutants, being putatively responsible for the observed impaired growth.

**ONE-SENTENCE SUMMARY:** Impaired pyrimidine nucleotide synthesis results in nucleotide limitation and imbalance, resulting in impaired photosynthesis, reduced growth, reproduction, and seed yield together with massively altered gene expression

## INTRODUCTION

Pyrimidine nucleotides are essential components of all living cells. They serve as building blocks for DNA and RNA and participate in metabolic processes ranging from sugar interconversion and polysaccharide metabolism to biosynthesis of glycoproteins and phospholipids (Kafer et al., 2004; Garavito et al., 2015). Most of nucleotides are incorporated into ribosomal RNA and thus influence translation and growth (Busche et al., 2020). Whereas the levels of free nucleotides are kept constant and balanced between purines and pyrimidines, ribosomal RNA pools dynamically respond to growth signals and during acclimation to cold (Busche et al., 2020, Garcia Molina et al., 2021). This response in RNA synthesis is controlled by target of rapamycin (TOR) i.a. by inducing expression of *de novo* synthesis genes aspartate transcarbamoylase (ATC) and dihydroorotate dehydrogenase (DHODH). Conversely, limiting nucleotide availability negatively affects TOR activity (Busche et al., 2020). Nucleotide metabolism can therefore be regarded as dynamic metabolic checkpoint and not as a static background process. In line with this, dramatic effects of pyrimidine nucleotide shortage on plastid DNA synthesis and photosynthetic performance have been observed in mutants with impaired activity of CTP synthase (Alamdari et al., 2021; Bellin et al., 2021b).

Pyrimidine biosynthesis is an ancient and evolutionarily conserved biochemical pathway and has been studied intensively in mammalian systems, other eukaryotes, and prokaryotes. Yet, studies of this pathway in plants are scarce, especially with respect to its regulation and interactions with other pathways. The first pyrimidine nucleotide, uridine monophosphate (UMP), is synthesized by the *de novo* pathway via enzymatic steps that appear to be invariant in all organisms (Martinussen et al., 2011). In most multicellular eukaryotes, including mammals, some fungi, and insects, the first three steps of the *de novo* pathway are encoded by a single transcriptional unit generating a polyprotein called CAD (Christopherson and Szabados, 1997; Kim et al., 1992; (Del Cano-Ochoa and Ramon-Maiques, 2021; Moreno-Morcillo *et al*., 2017). The CAD complex consists of carbamoyl phosphate synthase (CPS), ATC and dihydroorotase (DHO) and localizes to the cytosol.

Plant *de novo* pyrimidine biosynthesis follows a distinct gene and cell compartment organization scheme relative to other organisms (Nara et al., 2000; Santoso and Thornburg, 1998). The first three enzymes are encoded by individual and unlinked genes (Williamson and Slocum, 1994; Williamson et al., 1996; Nara et al., 2000) and the encoded proteins also exhibit different subcellular localizations (Witz et al., 2012; Witte and Herde, 2020). Although the first step in the pathway is encoded by CPS, ATC is responsible for the first committed step in plant pyrimidine biosynthesis. It localizes to the chloroplast stroma and catalyzes the production of carbamoyl aspartate (CA), which is then likely exported to the cytosol and converted to dihydroorotate by the cytosolic DHO enzyme.

Evidence is mounting for an association between the ATC and DHO enzymes at the chloroplast membrane (Doremus and Jagendorf, 1985; Witte and Herde, 2020; Trentmann et al., 2020; Bellin et al., 2021a), which would allow for metabolite channeling across cellular compartments. The localization of ATC in the chloroplast brings along the need for organellar import of ATC and the export of the enzyme product CA to the cytosol. This complication is accompanied by unique features of plant ATC. Plant ATC proteins are simple in structure, only consisting of a homotrimer as functional unit. Due to mutations in the active site, the homotrimer is under allosteric control and uniquely feedback inhibited by uridine monophosphate (UMP) (Bellin et al., 2021a). We think the simple structure is beneficial for fast import and assembly in the chloroplast whereas feedback inhibition is required for fine-tuning of ATC activity. DHODH, which resides in the mitochondrial intermembrane space, where it is coupled to the respiratory chain facilitates the closure of the pyrimidine ring to generate orotate (Zrenner et al., 2006; Witz et al., 2012). DHODH from multicellular eukaryotes, including Arabidopsis, requires ubiquinone as electron acceptor for activity, provided by the mitochondrial respiratory chain. Besides exhibiting a mitochondrial targeting peptide at its N terminus, Arabidopsis DHODH contains a transmembrane helix that anchors the protein into the inner mitochondrial membrane (Ullrich et al. 2002; Löffler et al., 2020). The last two enzymatic steps leading to the production of the first nucleotide, UMP, are catalyzed by the bifunctional cytosolic protein uridine-5′-monophosphate synthase (UMPS) (Nasr et al., 1994; Zrenner et al., 2006). The reason behind the distinct localization of each enzyme, in particular the chloroplast localization of ATC, is currently unclear.

RNA interference (RNAi) was previously utilized to knock down the expression of *ATC* in Arabidopsis and *ATC* and *DHODH* in Solanaceous species (Schröder et al., 2005; Chen and Slocum, 2008). A clear growth limitation was observed when *ATC* transcript levels were reduced by at least 50%, or 90% for *DHODH* (Schröder et al., 2005). Similarly, impaired growth of *ATC ami-RNA* lines but better growth in corresponding overexpressor lines was observed by Bellin et al., (2021), indicating that ATC is not present in large excess in Arabidopsis. In this work we aim to unravel further functions of the organellar located enzymes ATC and DHODH by in depth analysis of corresponding knock-down mutants. We observed most marked effects for ATC mutants in metabolism, gene expression, chloroplast function and ultrastructure. Although DHODH mutants were less affected, unique responses were identified as well.

## Results

### ATC mutants were most markedly impaired in growth and pyrimidine levels

We recently provided a preliminary characterization of a set of ATC mutants which contained two lines denominated as *atc#1* and *atc#2* which contained 17% and 12% residual amounts of transcript leading to 34% and 12% residual protein, respectively (Bellin et al., 2021a). This study was now complemented by analysis of two additional knock-down lines for DHODH, denominated as *dhodh#1* and *dhodh#2* exhibiting a residual transcript accumulation of 18% and 6%, respectively (Figure 1A). It was an obvious observation that *ATC* mutants were more severely impaired in growth and development compared to *DHODH* mutants. In comparison to the control plants (Col-0) all mutant lines showed a delay in the emergence of the first true leaves (Figure 1B, C). These early delays in development were followed by retarded and reduced growth in the vegetative and reproductive growth phase (Figure 1B, C) (Bellin et al., 2021) resulting in lower fresh weights (Figure 1D, E) (Bellin et al., 2021a) and smaller rosette diameters (Supplemental Figure S1A). Chlorosis was observed in *ATC* mutants only and accordingly chlorophyll levels were reduced in these lines by 41% and 64% (Figure 1F). In addition, the *ATC* mutant lines exhibited a significant reduction in the maximal primary stem length. Col-0 plants reached a maximum height of 40.5 cm, *atc#1* and *atc#2* only reached a height of 15.7 and 4.4 cm and *dhodh#1* and *dhodh#2* of 24.8 cm and 8.1 cm respectively (Supplemental Figure S1B, C).

**Figure 1.**
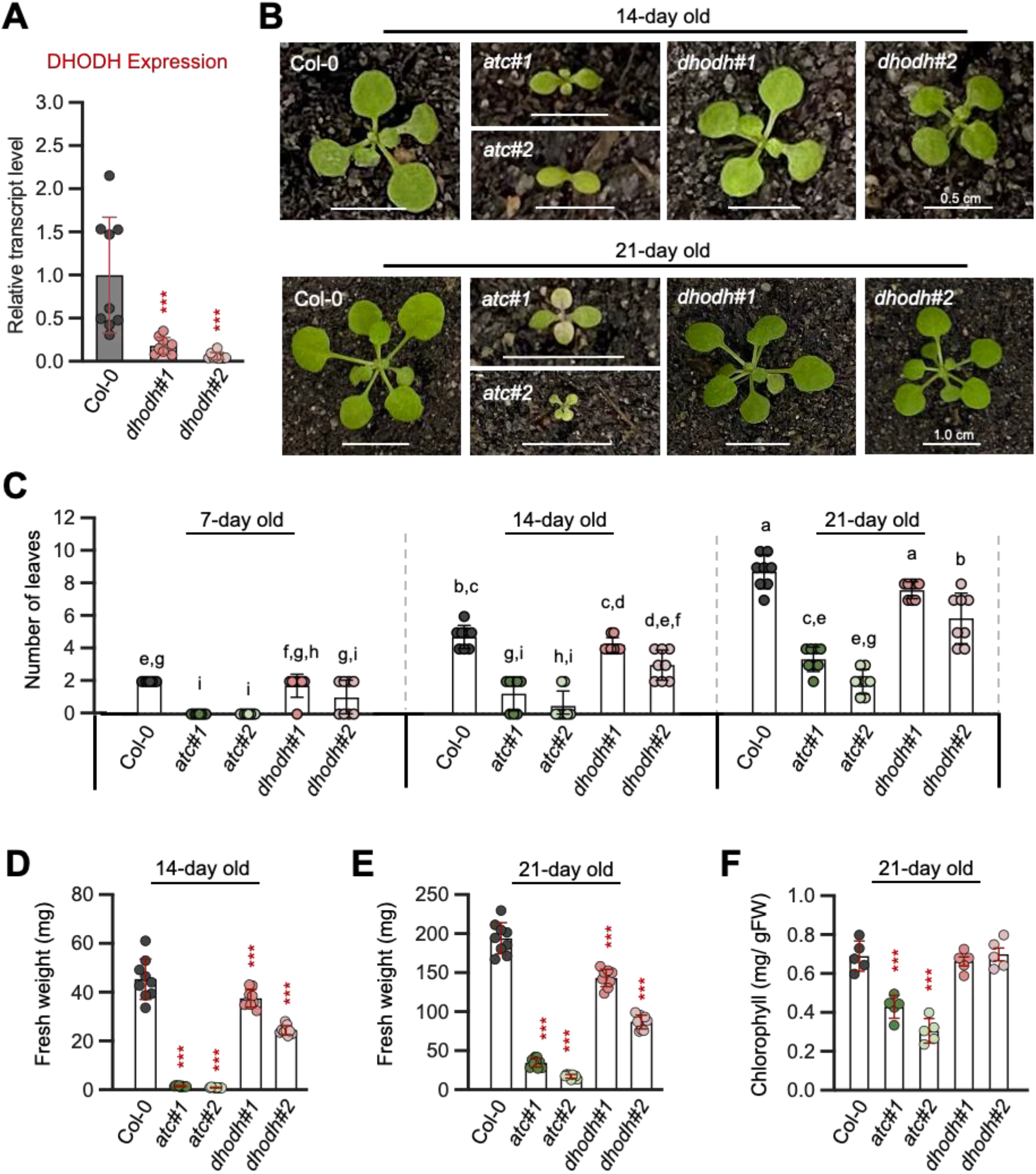
Silencing *ATC* and *DHODH* inhibits vegetative and reproductive growth. (**A**) *DHODH* transcript levels in *DHODH* knockdown mutants relative to Col-0. Expression was normalized to actin as a reference gene (n = 9). (**B**) Representative plants of 14- and 21-days old Col-0, *ATC* (*atc#1 & atc#2*) and *DHODH* (*dhodh#1* & *dhodh#2*) knockdown plants. (**C**) Number of leaves were counted after 7, 14 and 21 days of growth. (**D, E**) Fresh weight and (**F**) chlorophyll contents were determined after 14 and 21 days of growth. Shown are the means of five biological replicates +/-SD. For statistical analysis in A, D, E, F One way ANOVA was performed followed by Dunnett’s multiple comparison tests (*** = p < 0.001). Different letters in C denote significant differences according to two-way ANOVA with post-hoc Turkey HSD testing (p < 0.5).

### Downregulation of ATC results in pyrimidine nucleotide limitation

In an untargeted metabolic profiling of central metabolites using high resolution mass spectrometry, six-week-old plants grown under long day conditions on soil were inspected. Among pyrimidines, the two intermediates produced by ATC (carbamoyl-aspartate) and dihydroorotase (dihydroorotate), were barely detectable in both *ATC* lines whereas *DHODH* knock-down lines showed wild-type-like levels. This pattern was congruent with the levels of the downstream generated metabolites UMP, UDP, UDP-Glc and UDP-GlcNac (Figure 2A, B). In both *ATC* knock-down lines the levels of UMP and UDP were significantly reduced to around 50% compared to Col-0 controls. Furthermore, it is well known that pyrimidine and purine metabolites must be balanced to support nucleotide and nucleic acid synthesis (Reichard, 1988). Among purine nucleotides, the abundances of AMP and GMP were markedly increased in *atc#1, atc#2*, and *dhodh#2*, thus exhibiting a negative correlation to the pyrimidine nucleotides described above. In contrast, ADP and ATP levels were only marginally affected in all lines. Massively reduced levels of NADH and NADPH in both ATC lines were observed in addition (Figure 2B).

**Figure 2.**
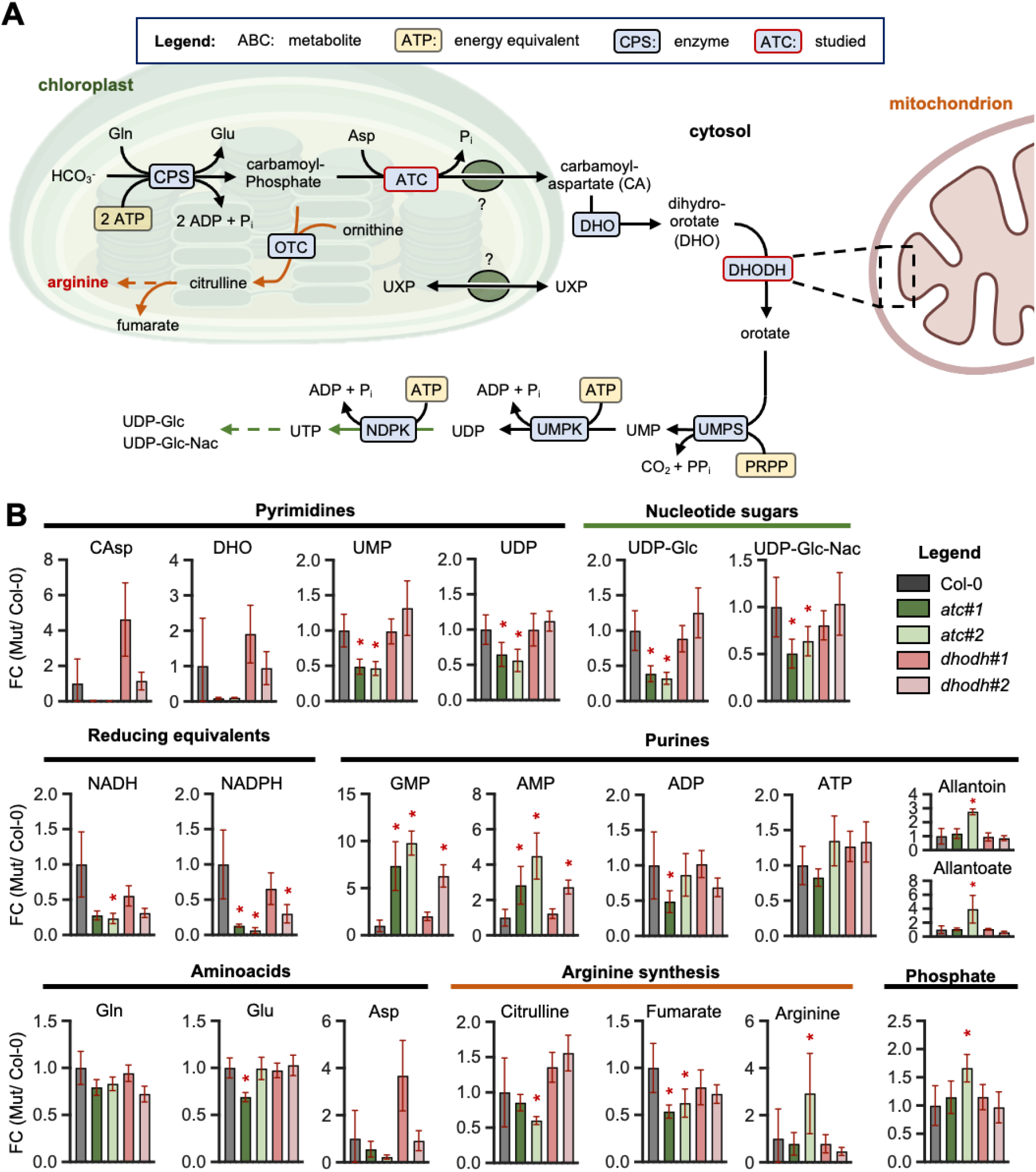
Scheme of pyrimidine de novo synthesis and corresponding metabolites levels. (**A**) Scheme of de novo pyrimidine biosynthesis pathway (black lines), arginine synthesis (orange lines) and biosynthesis of nucleotide sugars (green lines). All involved enzymes are highlighted in blue and deriving energy equivalents in yellow. ATC, aspartate transcarbamoylase; DHO, dihydroorotase; DHODH, dihydroorotate dehydrogenase; UMPS, UMP synthetase; CPS, carbamoyl phosphate synthetase; OTC, ornithine transcarbamoylase; UMPK, UMP kinase; NDPK, Nucleotide diphosphatekinase. (**B**) Relative metabolite levels from fully developed leaves are shown. Metabolite levels are shown as fold change (FC) relative to Col-0, which was set to 1. Data points represent means of five biological replicates ± SD. For determination of statistical significance (p-value < 0.05) Wilcoxon Mann-Whitney U-test was performed. Asterisks indicate significantly altered levels compared to Col-0.

Especially *atc#2* exhibits increased levels of the purine breakdown products 2-ureido glycine, allantoate, and allantoin, the latter are suggested to function in attenuating ROS stress (Brychkova et al., 2008). Several glucosinolates (glucoraphanin, glucoiberin, glucohirsutin, glucoiberin and glucohesperin), involved in pathogen resistance and constituting highly abundant secondary metabolites in Brassicaceae, were also strongly reduced in *atc#1, atc#2*, and *dhodh#2* (Supplemental Figure S2). Moreover, the levels of several intermediates of sugar metabolism, (glycolysis and TCA cycle) were significantly altered in the knock-down lines (Supplemental Figure S2).

### Transcriptome analysis revealed massively changed gene expression

Metabolic changes are mostly based on strong changes at the transcriptional level. To investigate which transcriptional changes occurred in the ATC mutant *atc#1*, global analyses were performed using RNA-Seq and compared with the corresponding control plants (Col-0). Therefore, overground tissue of six-week-old plants was harvested, RNA extracted and processed by standard RNA-Seq protocols at Novogene (China, UK). Compared to Col-0, a total of 2757 differentially expressed genes (DEGs) were significantly altered (p_adj._ < 0.05) in *atc#1* knockdown mutant. More detailed studies showed that 1100 genes (FC < 0) had moderate reductions (light blue) and 301 DEGs with an FC < -1 had significantly greater reductions in transcript levels in *atc#1* knockdown mutant (dark blue) (Figure 3A, B). In contrast, 1026 DEGs showed slightly (FC > 0) increased amounts and 330 other DEGs showed strongly increased transcript amounts (FC > +1) (Figure 3A, B). To determine which metabolic pathways were affected, Gene Onthology (GO) enrichment analysis was performed. Listed are selected biological processes which were most significantly altered (-Log_10_p_adj._) (Figure 3C, D). Polysaccharide metabolic processes and cell wall organization had the largest number of reduced expressed genes in *atc#1* knockdown mutant compared to Col-0 with 92/436 and 92/471, respectively (Figure 3C). Other affected metabolic pathways with reduced transcripts includes are the generation of precursor metabolites and energy (81/482), nucleobase-containing small molecule metabolic process (76/450), response to metal ions (74/475), photosynthesis (51/273), sulfur compound metabolic process (60/388) and the cellular amino acid metabolic process (60/460) (Figure 3C). Upregulated genes belong to the GO terms RNA splicing (46/304), mRNA processing (57/433), as well as plastid organization (43/280). ribonucleoprotein complex biogenesis (62/490) processing of proteins to the chloroplast (14/49), plastid transcription (6/11), protein refolding (13/49), rhythmic process (23/149), response to oxidative stress (49/451) and the hydrogen peroxide metabolic process (16/99) (Figure 3D).

**Figure 3.**
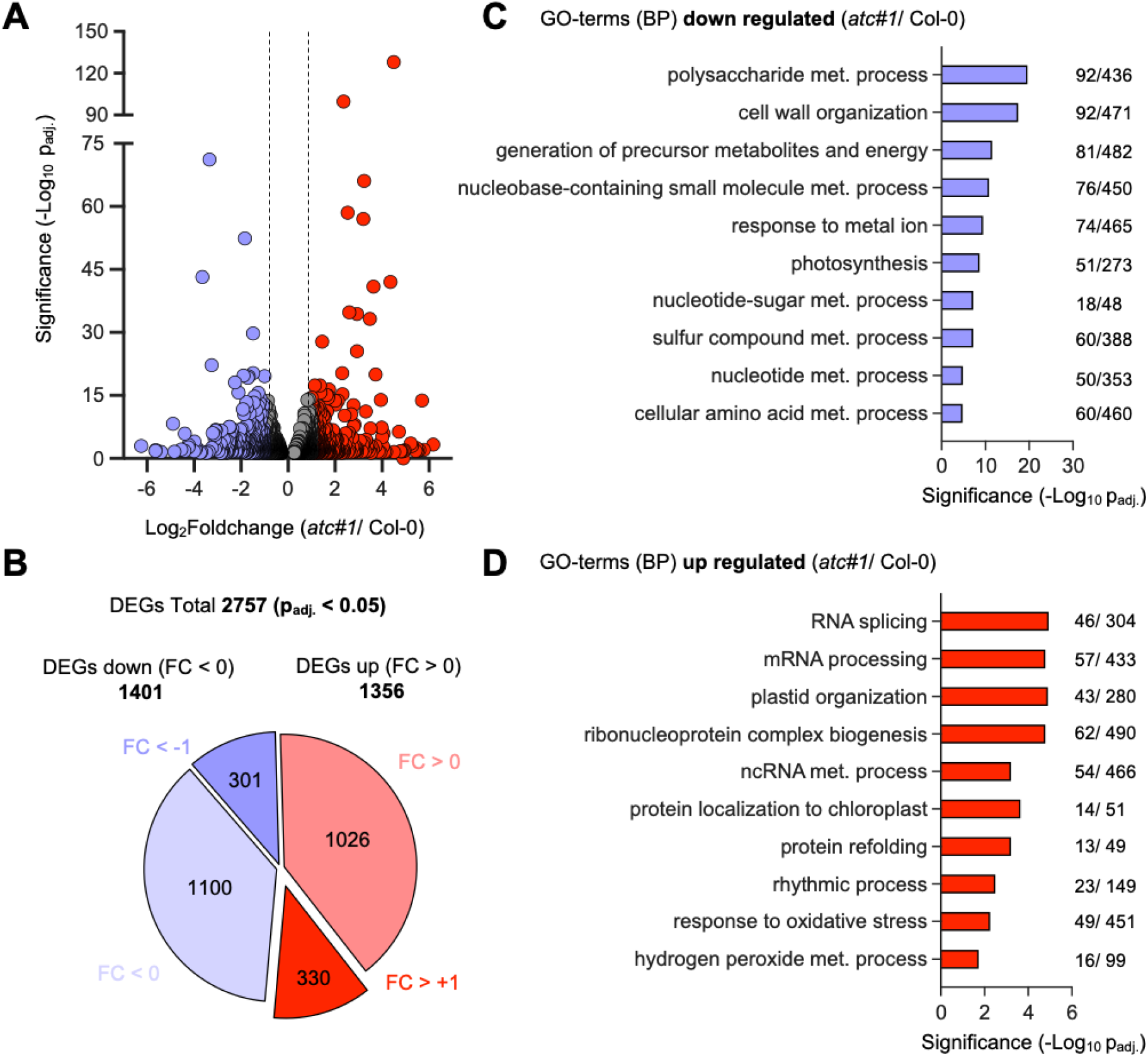
RNA Seq analyses of the transcriptome from leaf tissue. The (**A**) volcano plot and (**B**) pie charts show significantly (p_adj._ < 0.05) differentially expressed genes (DEGs) in *atc#1* knockdown mutant compared to Col-0. Changes in expression are shown as Log_2_Foldchange. Reduced expression was detected for 1401 genes (blue) and increased expression for 1356 genes (red). Thereby 301 DEGs showed a Log_2_FC < -1 and 330 DEGs a Log_2_FC > +1. Detected DEGs were subdivided by GO-terms analysis into different biological processes (BP) that were either (**C**) enriched or (**D**) repressed in *atc#1* mutants compared to Col-0.

More detailed studies revealed reduced expression in the groups “polysaccharide metabolic processes”, “cell wall organization”, and “photosynthesis” (Supplemental Figure S3A). Prominent upregulated pathways were “oxidative stress response”, “circadian rhythm”, and “ribonucleoprotein complex synthesis” (Supplement Figure S3B, C). DEGs in the pathway of nucleotide metabolism revealed downregulation of purine *de novo* synthesis with reduced expression of ADSL. However, IMPDH and GMPS leading to GMP synthesis were increased. Genes of salvage pathway enzymes showed reduced expression except for the plastidic NDPK2 (Table 1). Genes of purine and pyrimidine catabolism were all reduced in expression (Table 1). When inspecting genes for intracellular metabolite transporters (based on the selection in (Linka and Weber, 2010), 14 DEGs were identified (Table 2). Interestingly, plastid localized NTT1 and 2 (ATP & ADP) transporters and the phosphate carrier PHT4.5 sowed increased expression, all other genes in this category were reduced in expression, including mitochondrial dicarboxylate carriers DTC and DIC, uncoupling protein UPC and the ATP/ADP translocator AAC1. (Table 2). Because metabolite levels pointed to altered energy metabolism, we checked for DEGs in carbohydrate (glycolysis and TCA cycle) and respiration. From 46 altered genes, only six showed increased expression. All these do not belong to the canonical players in the respective pathways but exert different functions, e.g. GAPN catalyzes a bypass reaction in glycolysis to serve mannitol production (Kirch *et al*., 2004), HKL, serving as a negative growth regulator (Karve and Moore, 2009) and alternative NADPH dehydrogenase and alternative oxidase (AOX1) in mitochondrial respiration not coupled to ATPproduction (Table 3). Marked reduction was observed for PDC3 (Log_2_FC -2.8), one of the two main pyruvate dehydrogenase complexes and cytosolic fumarase 2 (FUM2, Log_2_FC-1.49) involved in carbohydrate partitioning and adaptation to abiotic stress, for example cold stress, (Dyson *et al*., 2016; Pracharoenwattana *et al*., 2010).

**Table 1.** DEGs in nucleotide metabolism. Manually selected genes in nucleotide metabolism are shown (FDR <0.05).

| Gene ID | Symbol or function | *atc#1* vs Col-0<br>(log 2fc) |
| --- | --- | --- |
| At3g20330 | ATC | **-3.34** |
| At2g37690 | AIRCAR | **0.69** |
| At4g18440 | ADSL | **-1.30** |
| At1g79470 | IMPDH1 | **0.76** |
| At1g16350 | IMPDH2 | **0.51** |
| At1g63660 | GMPS | **0.73** |
| At3g23580 | RNR2A | **0.53** |
| At1g72040 | DNK | **0.63** |
| At5g50370 | AMK3 | **-0.60** |
| At5g26667 | UMK3 | **-0.69** |
| At5g63310 | NDPK2 | **0.41** |
| At4g11010 | NDPK3 | **-0.71** |
| At4g22570 | APRT3 | **-0.74** |
| At3g09820 | ADK1 | **-0.68** |
| At5g03300 | ADK2 | **-0.47** |
| At5g28050 | GSDA | **-0.52** |
| At3g17810 | PYD1 | **-0.53** |
| At5g12200 | PYD2 | **-0.41** |
| At5g64370 | PYD3 | **-0.41** |
| At2g26230 | UOX | **-0.37** |

**Table 2.** DEGs in central carbon metabolism. Genes selected from the KEGG database with functions in central carbon metabolism and respiration are shown (FDR <0.05).

| Gene ID | Symbol or (function) | atc#1 vs Col-0<br>(log 2fc) |
| --- | --- | --- |
| At3g03250 | UGP | -0.57 |
| At1g23190 | PGM3 | -0.53 |
| At1g20950 | PFP-ALPHA1 | -0.55 |
| At3g55440 | CTIMC | -0.40 |
| At2g24270 | ALDH11A3 | 0.30 |
| At4g26520 | (Aldolase) | 0.40 |
| At1g53310 | PPC1 | -0.65 |
| At1g70280 | (PGM) | 0.52 |
| At2g22480 | PFK5 | -0.43 |
| At5g56350 | (Pyruvate Kinase) | -0.46 |
| At4g19130 | HXK1/ GIN2 | -0.34 |
| At1g50460 | HKL1 | 0.53 |
| At1g66430 | FRK3 | -0.98 |
| At5g51830 | FRK1 | -0.53 |
| At5g56630 | PFK7 | -0.35 |
| At3g52930 | FBA8 | -0.38 |
| At5g01330 | PDC3 | -2.80 |
| At1g10670 | ACLA-1 | -0.42 |
| At4g35260 | IDH1 | -0.51 |
| At3g55410 | (2-oxoglutarate DH) | -0.33 |
| At5g55070 | (2-oxoglutarate DH) | -0.62 |
| At5g65750 | (2-oxoglutarate DH) | -0.43 |
| At2g20420 | (Succinate-CoA ligase) | -0.35 |
| At5g08300 | (Succinyl-CoA ligase) | -0.49 |
| At3g27380 | SDH2-1 | -0.43 |
| At5g40650 | SDH2-2 | -0.66 |
| At5g50950 | FUM2 | -1.49 |
| At2g47510 | FUM1 | -0.31 |
| At3g15020 | mMDH2 | -0.58 |
| At1g04410 | MDH1 | -0.32 |
| At5g43330 | MDH2 | -0.78 |
| At2g22780 | PMDH1 | -0.32 |
| At5g47890 | (NADH Ubi-<br>oxidoreductase) | -0.37 |
| At1g07180 | NDA1 | 0.59 |
| At5g43430 | ETFBETA | -0.50 |
| At1g36380 | (Cytochrome-c reductase) | -0.77 |
| At1g15120 | (Cytochrome-c reductase) | -0.44 |
| At5g13430 | UCR1-1 | -0.36 |
| At4g32470 | QCR7-1 | -0.60 |
| At5g08670 | (ATP synthase- beta SU) | -0.28 |
| At5g13450 | ATP5 | -0.30 |
| At1g51650 | (ATP synthase- delta SU) | -0.48 |
| At3g52300 | ATPQ | -0.47 |
| At5g47030 | (mATP synthase- deltaS U) | -0.55 |
| At2g33040 | ATPC | -0.49 |
| At3g22370 | AOX1A | 0.70 |

**Table 3.**
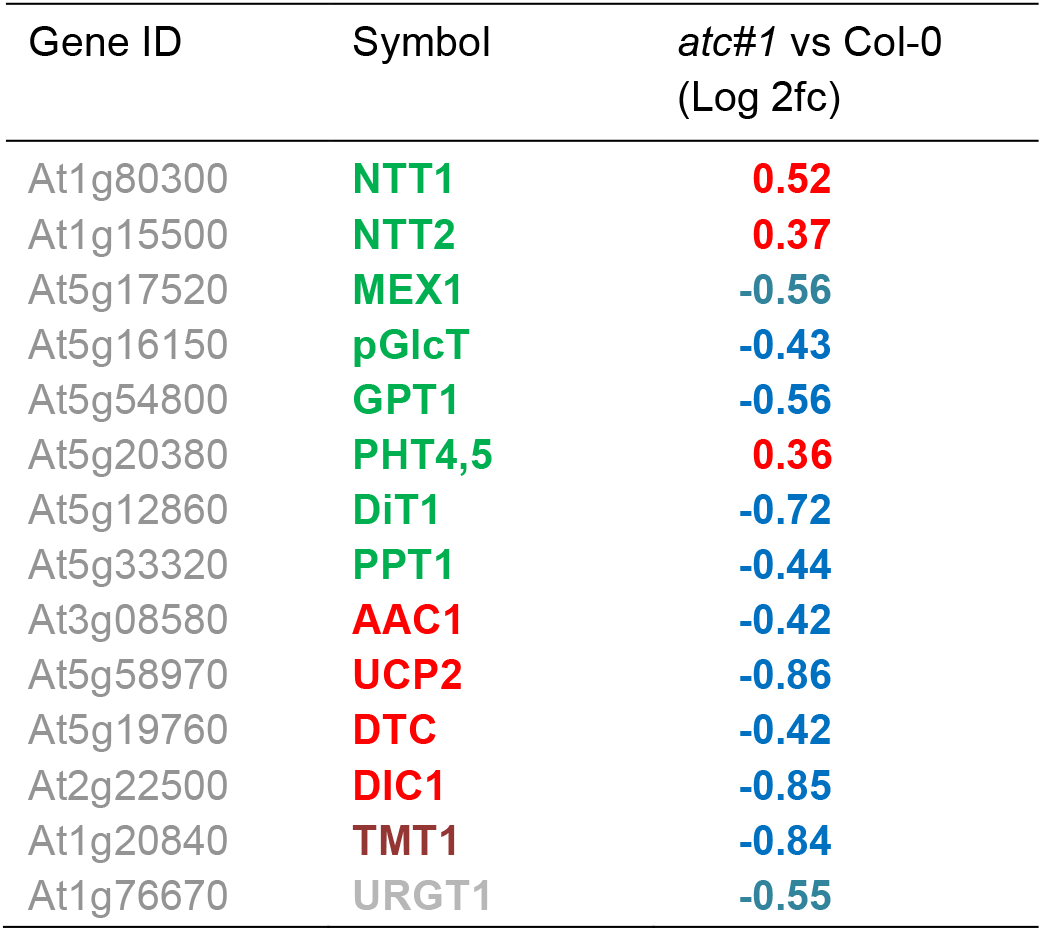
DEGs in intracellular metabolite transport. Transport proteins active in intracellular metabolite transport were extracted from the literature and DEGs with FDR < 0.05 are shown. Color of symbols: green = plastid localization, red = mitochondrial localization, black = vacuolar localization, grey = golgi localization.

### Tissue specific expression of ATC and DHODH are similar

To examine whether the observed differences between *ATC* and *DHODH* mutants can be attributed to differences in tissue-specific expression, transgenic plants were produced. Typical examples of GUS staining patterns are shown in Supplemental Figure S4. In all experiments, *ATC::GUS* and *DHODH::GUS* reporter constructs exhibited similar expression patterns. A developmental time course revealed that *ATC* (Supplemental Figure S4 A-H) and *DHODH* (Supplemental Figure S4I-P) are highly expressed during seed germination and early seedling development (Supplemental Figure S4 A-C and I-K). In two-week, old seedlings GUS signal could be detected over the whole cotyledons (Supplemental Figure 4C, K). Furthermore, the vasculature showed intense staining in cotyledons. Staining of the leaf vasculature remained high during leaf development, in contrast, staining in the mesophyll weakened with increasing age of the leaves (Supplemental Figure S5). Moreover, intensive staining was also visible in primary and secondary roots as well as in root tips (Supplemental Figure S4D, E, l, M).

### Embryo-, seed development are altered in knock-down lines in pyrimidine *de novo* synthesis

When developing siliques of mutant lines were inspected, empty positions with aborted seeds (red asterisks) and less colored seeds (lacking embryos; white arrows) were visible in all knock-down lines, but not in Col-0 controls (Figure 4A). The number of seeds per silique was found to be reduced in all knock-down lines, but strongest in *atc#2* with only 22.1% of residual viable seeds (Figure 4B).

**Figure 4.**
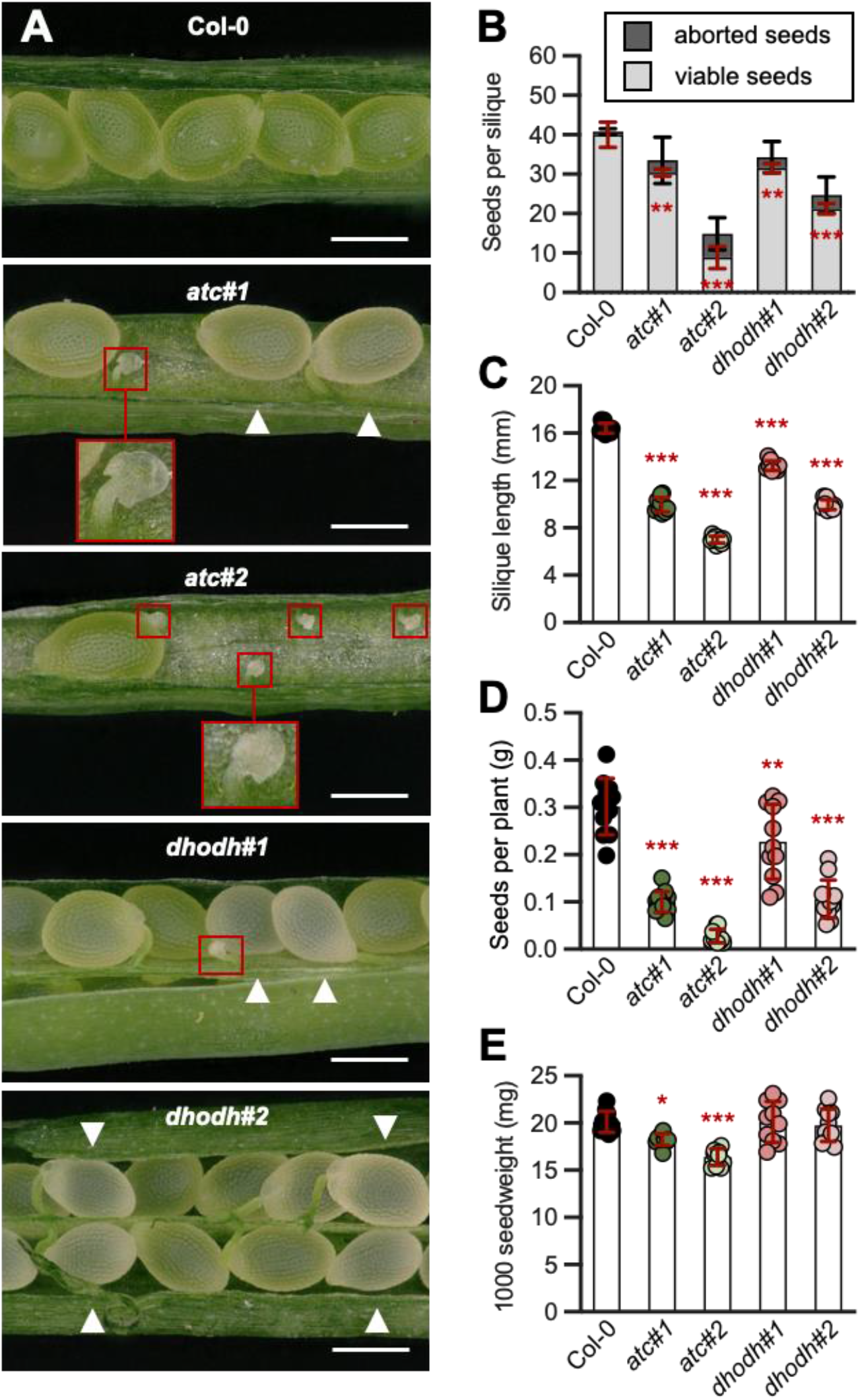
Embryo and seed development. (**A**) Representative siliques of Col-0, *ATC* and *DHODH* knock-down lines showing viable, aborted seeds (red box) and lacking embryos (white arrow). (**B**) Number of viable and aborted seeds per silique were counted from 10 siliques of five different plants per line. (**C**) To determine siliques length 10 siliques of 10 different plants per genotype were analyzed. (**D**) Seed weight per plant (n=11). (**E**) For determination of 1.000 seed weight 10 different plants per line were used. Data points represent means of biological replicates ± SD. Asterisks depict significant changes between the different lines referring to the WT according to one-way ANOVA followed by the Dunnett’s multiple comparison test (* = p < 0.05, ** = p <0.01, *** = p<0.001). Scale bar in A = 2 mm.

Silique length was reduced in all mutant lines. Compared to control plants the silique length in *ATC* knock-down lines was reduced to 60.5% and 42.6%, and for *DHODH* knock-down lines down to 80.7% and 60.7% (Figure 4C). Shorter siliques as well as increased numbers of aborted seeds per silique in knock-down lines resulted in reduced yield of mature seeds per plant. The weight of seeds per plant was reduced by 66% and 91% for *atc#1* and *atc#2* and by 24% and 65% for *dhodh#1* and *dhodh#2* in comparison to the Col-0 (Figure 4D). Analysis of the 1.000-seed weight of mature, dried seeds revealed a reduced seed weight in both *ATC* knock-down lines, whereas in *DHODH* knock-down plants the 1.000-seed weight was comparable to the wild type (Figure 4E).

To determine whether the seed development impacts mature seed properties, the seed germination was analyzed. Whereas only small alterations were observed between *Col-0* and *ATC* knock-down lines, surprisingly both *DHODH* knock-down lines showed a significant delay in germination (Figure 5A). 30 hours after transfer of seeds to ambient growth conditions, 97% of Col-0 and 93% and 94% of *atc#1* and *atc#2* seeds germinated. For *DHODH* knock-down lines only 44% (*dhodh#1*) and 30% (*dhodh#2*) of the seeds were germinated after the same time (Figure 5A). Rescue experiments with uridine and uracil (1mM each) did not support germination in DHODH mutants, but uracil provoked delayed germination in Col-0 and ATC mutants (Figure 5B, C).

**Figure 5.**
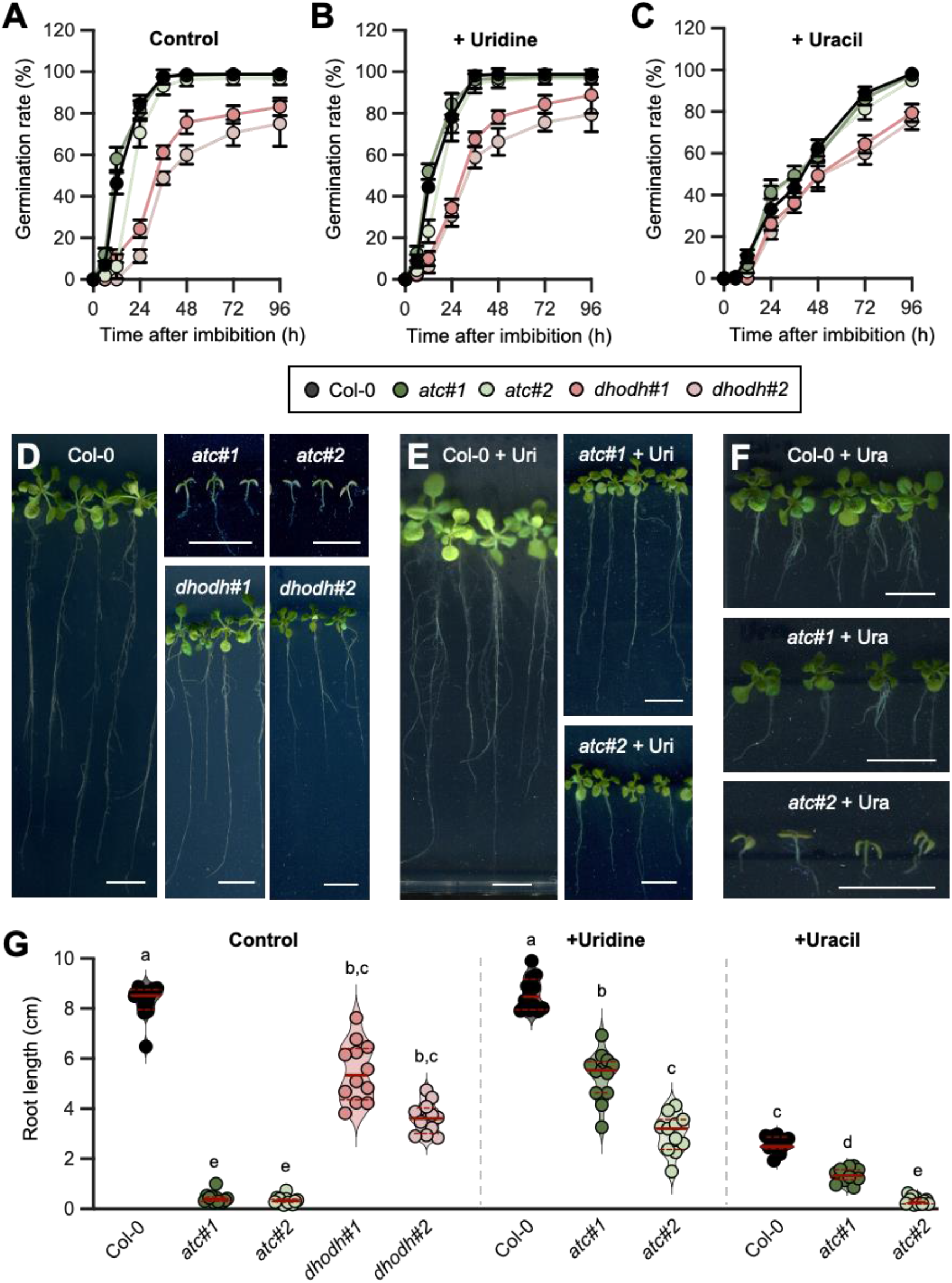
Seed development and supplementation studies. (**A-C**), Germination rate monitored in a time course of 48 hours after imbibition. For (**A**) control conditions plants were grown on ½ MS-Medium without any supplementation in a 14h light/10h dark regime. To determine the effect of (**B**) uridine or (**C**) uracil on seed germination ½ MS-Medium was supplemented either with 1 mM uridine or uracil. (**D-F**) Typical examples of 21 days old plants which were grown on (**D**) ½ MS-Media and supplemented with (**E**) uridine or (**F**) uracil. (**G**) Determination of root length shown in D-F. Data points represent means of biological replicates ± standarddeviation. For statistical analysis in A-C One way ANOVA was performed followed by Dunnett’s multiple comparison tests (*** = p < 0.001). Different letters in G denote significant differences according to two-way ANOVA with post-hoc Turkey HSD testing (p < 0.5). Scale bar in **D-F** = 1 cm.

Monitoring of root growth revealed that *DHODH* knock-down plants had compensated their germination delay within five days and appeared similar to wild-type plants, whereas the development of *atc#1* and *atc#2* was nearly arrested 5 days after germination (Figure 5D). Supplementation with uridine partially rescued growth delays in ATC mutants and uracil supported growth in *atc#1* but at the same time reduced growth in Col-0 (Figure 5E-G).

### Knock-down lines in pyrimidine *de novo* synthesis reveal altered ultrastructure of chloroplasts and mitochondria

Since all mutant lines showed a severely reduced growth, the leaf morphology and cellular ultrastructure were analyzed in detail by means of histology and transmission electron microscopy. For this, we focused on the most affected lines *atc#2* and *dhodh#2*. Light microscopy analysis of cross sections from mature leaves revealed that leaf thickness was reduced by approximately 26% and 5% in *atc#2* and *dhodh#2* compared to corresponding control plants (Figure 6A-F) (Supplemental Figure S6A). However, whereas the leaf architecture was wild-type like in *dhodh#2, atc#2* knock-down lines showed an altered architecture: the layer of palisade parenchyma was disturbed and the intercellular space in the spongy parenchyma was less pronounced. Furthermore, the number of chloroplasts was reduced in *atc#2* (Figure 6A-F).

**Figure 6.**
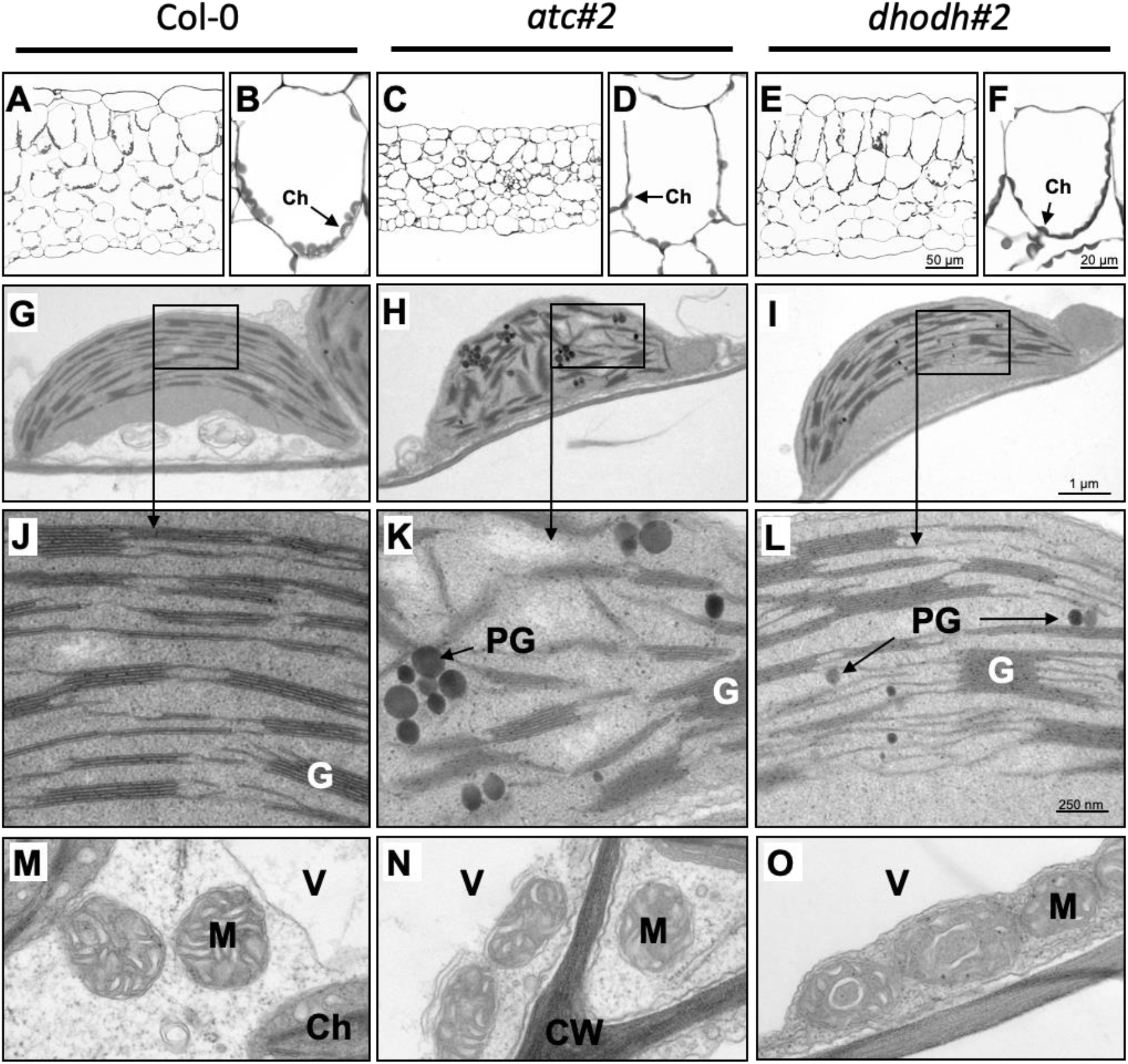
Histological and ultrastructural analysis of rosette leaves of Arabidopsis Col-0, *atc#2* and *dhodh#2*. Light (**A-F**) and transmission electron microscopy images (**G-O**) of leaf cross sections of Arabidopsis Col-0 (**A, B, G, J, M**), *atc#2* (**C, D, H, K, N**) and *dhodh#2* (**E, F, I, L, O**). Histological cross section of rosette leaves (**A, C, E**) with close up of a palisade parenchyma cell (**B, D, F**). Ultrastructure of chloroplasts with close ups of thylakoids (**G-L**) and mitochondria (**M-O**) with changes of ultrastructure partly to be observed in dhodh#2 plants (**O**, arrow heads). Ch, chloroplast; CW, cell wall; ER, endoplasmic reticulum; G, grana; M, mitochondria; PG, plastoglobuli; V, vacuole.

Transmission electron microscopy (TEM) revealed that chloroplasts of Col-0 (Figure 6G, J) and *dhodh#2* (Figure 6I, L) plants showed well-developed thylakoids, with typical stacked and interconnected grana. Chloroplasts of *atc#2* mutant plants exhibited changed ultra-structure characterized by loose appearing thylakoids, less dense stacked grana (Figure 6H, K). In addition, their arrangement within the chloroplast was significantly more irregular than in the wild type plants. Further analysis revealed that chloroplasts of *dhodh#2* plants show the same size as the Col-0, whereas the chloroplast size of *atc#2* of is reduced by about 35% (Supplemental Figure S6B).

Since the chloroplast ultrastructure was largely unchanged in *dhodh#2;* we intended to determine if the observed phenotype of *DHODH* knock-down lines might be based on defects in the mitochondrial ultrastructure. Thereby, no significant differences in mitochondrial sizes were observed between wild-type and mutant lines (Supplemental Figure S6C). However, compared to Col-0 and *atc#2* plants about 16% of the mitochondria from *dhodh#2* showed an altered ultrastructure in which the granules were less abundant, and the cristae formed by the inner membrane were reduced. Additionally, the formation of ring like structures has been observed (Figure 6O; black arrows).

### Altered photosynthetic efficiency in knock-down lines in pyrimidine *de novo* synthesis

To determine alterations in the physiology of the lines analyzed, which are fundamental for the observed growth and morphological alterations, we first measured PSII efficiency using chlorophyll fluorescence imaging in a light curve setting. Thereby, Col-0 and *dhodh#1* and *dhodh#2* plants exhibited almost identical maximal photosynthetic efficiency (0.8) whereas *atc#1* and *atc#2* plants showed a reduction to only 0.53 and 0.50, respectively (Figure 7A). To prevent photosystem II (PSII) from photodamage plants dissipate light energy as heat in the process of non-photochemical quenching (NPQ), thus lowering photosynthetic efficiency (Φ_PSII_) (Müller et al., 2001; Lambrev et al., 2012). This was reflected by reduced maximal Φ_PSII_ values and higher NPQ in *ATC* knock-down lines (Figure 7A). NPQ values for *dhodh#1* were close to those of control plants, while values of *dhodh#2* were intermediate between Col-0 and *ATC* knockdown line*s* (Figure 7A). As we speculate about an effect of reduced pyrimidine nucleotides on rRNA abundance and as related consequence impaired synthesis of photosynthesis related proteins, these were quantified by immunoblotting. The main proteins of photosynthetic reaction centers, PsaA, PsbA, and PsbD were found reduced in *atc#1* and *atc#2* compared to Col-0 and *DHODH* knock-down lines. PetC was not affected and AtpB was decreased in all mutant lines (Figure 7B).

**Figure 7.**
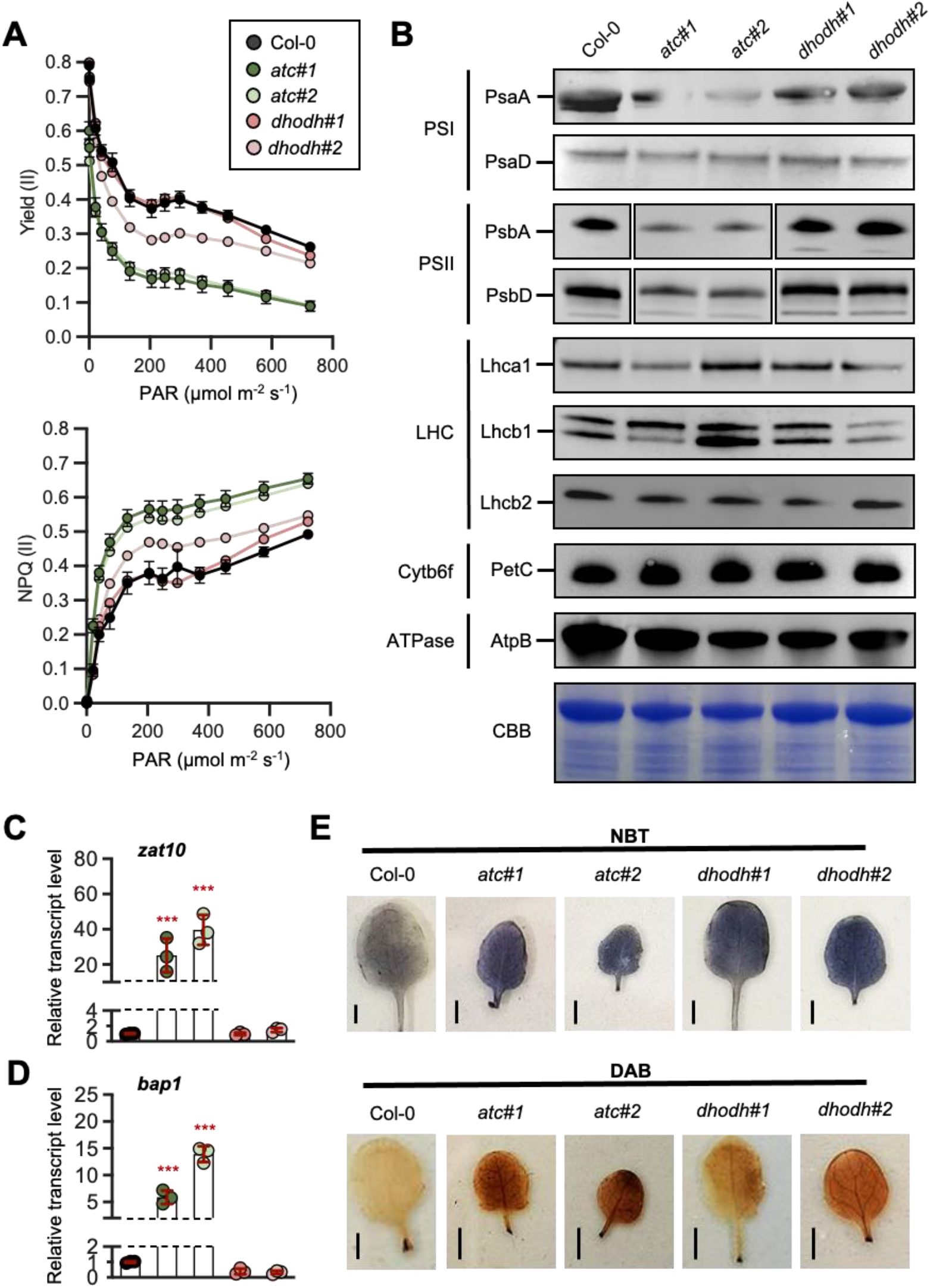
Determination of photosynthetic efficiency and ROS related parameters. (**A**) Photosynthetic efficiency of photosystem II (yield (II)) and non-photochemical quenching (NPQ (II)) of four weeks old Col-0, ATC- and DHODH-knockdown plants were measured in a light response curve (n ≥ 8). (**B**) Immunoblot analysis of photosynthesis related proteins. Proteins were extracted from leaves on a denaturing gel and probed with antibodies as indicated. CBB: Coomassie brilliant blue as a loading control. (**C, D**) Relative transcript levels of chloroplast ROS signalling markers (A) *zat10* (**D**) and *bap1* were normalized to actin and Col-0 was set to 1 (n = 3). (**E**) Accumulation of O_2_− and H_2_O_2_ in two-week-old plants grown under 14h light/ 10h dark regime visualized by NBT (top panel) and DAB staining (bottom panel) in leaves harvested after 6 hours of light. Images show typical results from the analysis of at least 5 leaves from 3 individually grown plants (Scale bar 0.25 cm). Data points are means ± (A) standard error and in (C, D) standard deviation. For statistical analysis in A-C One way ANOVA was performed followed by Dunnett’s multiple comparison tests (*** = p < 0.001).

### ROS accumulation challenges the detoxification system in *ATC* knock-down mutants

Impaired photosynthetic efficiency can be caused by increased reactive oxygen species (ROS) production (Kato et al, 2009, Su et al., 2018). Thus, the expression of two chloroplast ROS signaling marker genes *zat10* and *bap1* were quantified and found to be massively increased in both *ATC* lines and only slightly in *DHODH* lines (Figure 7C, D). Superoxide and H_2_O_2_ were visualized using nitroblue tetrazolium (NBT) and 3,3-diaminobenzidine (DAB; Figure 7E). NBT staining showed that superoxide accumulation was increased in leaves of all knock-down lines relative to Col-0 control plants (Figure 7E, top panel). DAB staining indicates that accumulation of H_2_O_2_ was strongly increased in *atc#1* and *atc#2*, whereas in the DHODH knock-down lines DAB staining was less intense (Figure 7E, lower panel).

### Altered assimilation and respiration in *atc#1* and *dhodh#1* knockdown mutants

Reduced photosynthetic efficiency came along with a reduced carbon assimilation rate (*A*) in *atc#1* (74%) and *dhodh#1* and (75%) of the wild type, respectively (Figure 7C). Respiration (*R)* decreased to 68% and 79% of wild-type level in *atc#1* and *dhodh#1* plant (Figure 7C).

## Discussion

In this work, we analyzed the function of two enzymes of the *de novo* pyrimidine biosynthesis pathway located in the chloroplast (ATC) and mitochondria (DHODH) by employing corresponding knock-down lines. *ATC* transcript levels were reduced by 84 and 90% relative to wild type in the two analyzed, representative knock-down lines, which corresponded to the lower levels of ATC protein (Bellin et al., 2021a) consistent with previous reports (Chen and Slocum, 2008). These lines showed severe growth limitations throughout development. Notably, *DHODH* lines had much weaker phenotypes than *ATC* lines, although both sets of lines were characterized by a comparable reduction in transcript levels (Figure 1A). There is no information available that would suggest different protein amounts or enzyme activity in response to transcript reduction in both mutants. However, it is not surprising that mutation in the first committed step of a pathway, here ATC, results in more pronounced phenotypes as mutations in later steps, because these might be compensated by upregulation of early steps, resulting in increased substrate availability or the possibility that compensatory salvage reactions rescue phenotypes. In support of this view, previous reports on mutants in *de novo* pyrimidine synthesis from Arabidopsis or solanaceous species, identified more pronounced phenotypic alterations in ATC mutants, compared to those encoding later pathway reactions. (Schröder et al., 2005; Geigenberger et al., 2005; Chen et al., 2008). Moreover, downregulation of UMPS even provoked an overcompensation by the salvage pathway, resulting in increased biomass accumulation. (Geigenberger et al., 2005).

The expression pattern of the *ATC* and *DHODH* genes did not reveal substantial differences when interrogated with promoter GUS reporter constructs. Both *ATC* and *DHODH* are expressed throughout development in roots, shoots and flowers, in line with previous reports on *ATC*-promoter-GUS studies in Arabidopsis (Chen and Slocum, 2008) or by Northern blots with probes for *de novo* pyrimidine synthesis genes in tobacco (*Nicotiana tabacum*) leaves (Giermann et al., 2002). This high expression throughout development indicates a constant substantial function of the pathway in Arabidopsis. However, reduced expression of *ATC* and *DHODH* in leaf mesophyll cells of older leaves was observed, suggesting a switch to pyrimidine salvage during aging, largely achieved via uridine/cytidine kinases, while expression remained high in leaf veins (Ohler et al., 2019).

Nucleotide limitation and pyrimidine/purine imbalances lead to severe problems in cell division and are causative of diseases in animals (Del Cano-Ochoa and Ramon-Maiques, 2021; Diehl *et al*., 2022) This aspect is markedly understudied in the plant field. However, in our work we observe a pyrimidine nucleotide limitation, going along with altered levels of purines (Figure 2) and a massive reprogramming of metabolism including pathways like nucleotide metabolism, intracellular transport and carbohydrate and energy metabolism (Figure 3, Tables 1-3).

Reduced amounts of various pyrimidine nucleotides (UMP, UDP, UDP-Glc and UDP-Glc-Nac) are accompanied by increased amounts of purine monophosphates (GMP and AMP) and indicate a low energy state in ATC mutants (Figure 2). Reduced expression of ADSL and increased levels of the breakdown products allantoin and allantoate indicate an attempt of the plant to balance nucleotide levels. Here, a general downregulation of purine *de novo* synthesis at the level of ADSL is accompanied by increases in IMPDH and GMPS, leading to GMP synthesis. In fact, GMP levels increase most strongly among all metabolites measured, up to 10-fold (Figure 2). Reduced expression of all three pyrimidine catabolic enzymes (PYD1-3) might reflect an attempt to stabilize pyrimidine levels. Reduced expression of GSDA and UOX, both acting in purine catabolism (Dahncke and Witte, 2013), might explain high GMP levels, but are counterintuitive of alleviated Allantoin and Allantoate (Figure 2). Although it’s unclear how the increase of both purine metabolites is achieved, they might function in reducing oxidative stress (Brychkova *et al*., 2008).

Nucleotide limitation apparently leads to reduced expression of genes in polysaccharide metabolism and cell wall organization as revealed by the GO term analysis (Figure 3) (Supplemental Figure S3). Changes in cell wall synthesis can probably be explained by the fact that a central metabolite, UDP-Glc, is strongly reduced. This might also explain reduced leaf thickness in *atc#2* knockdown mutants (Figure 6) (Supplemental Figure S6).

Other observed effects in *ATC* mutants were related to altered regulation of central carbohydrate metabolism and mitochondrial energy generation. Most of the corresponding genes in glycolysis, TCA cycle, respiration and corresponding metabolite transporters showed downregulation (Table 2, Table 3). This is reflected by reduced amounts of isocitrate in *atc#1*, and succinate and fumarate in *atc#1* and *atc#2* (Figure 2, Supplemental Figure S2). Major alterations in expression occurred with Pyruvate Decarboxylase 1 (PDC3, -2.8) and Fumarase 2 (FUM2, -1.49) (Table 2). PDC3 marks the entry point of carbohydrates to TCA cycle and cytosolic FUM2 is involved in nitrogen assimilation and growth under high nitrogen (Pracharoenwattana et al., 2010). In addition, FUM2 plays a role in cold acclimation (Dyson et al., 2016). FUM2 knockouts show reduced fumarate levels but increased malate, as observed in ATC mutants as well (Figure 2, Supplemental Figure S3). Increased levels of hexose phosphates indicate that not substrate limitation leads to suppression of central carbohydrate metabolism. Interestingly, both plastidic ATP/ADP carriers are upregulated, supposedly to supply the chloroplast with extra energy, while the mitochondrial ATP/ADP carrier ACC1 is downregulated (Table 3). Reduced photosynthetic capacity reflected in reduced assimilation seems to be balanced by reducing respiration (Table 4).

**Table 4.**
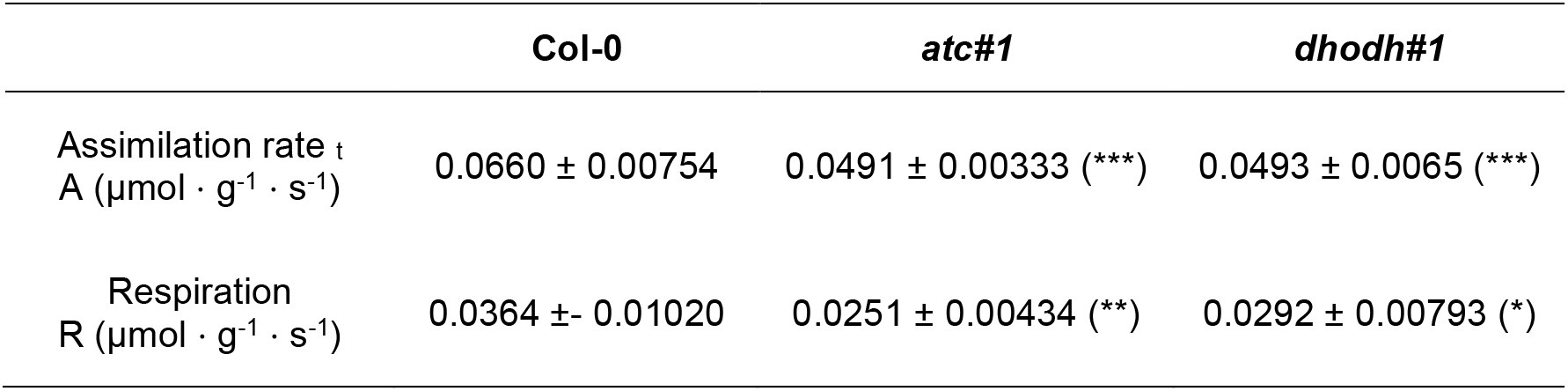
CO_2_-Assimilation- and Respiration rate. Determination of assimilation and respiration rate by Gas-exchange measurements. Plotted are mean values of eight biological replicates ± standard error. For statistical analysis one way ANOVA was performed followed by Dunnett’s multiple comparison tests (* = p < 0.05, ** = p < 0.005, *** = p < 0.001).

The observed massive change in gene expression in *ATC* lines is surely driven by a central regulatory process. The Target of Rapamycin (TOR) complex represents such a central growth regulator in plants and animals.

Growth requires the constant biosynthesis of ribosomes, which will place a significant demand for nucleotides to provide ribosomal RNA which can consume up to 50% of all cellular nucleotides (Busche et al., 2020). Among the pyrimidine biosynthesis genes, *ATC* and *DHODH* were shown to be upregulated by the glucose-TOR complex (Xiong et al., 2013). Conversely, nucleotide limitation negatively affected TOR activity. It is thus likely that nucleotide limitation in ATC mutants, but not in DHODH causes the large reprograming of metabolism via altered gene expression, partially regulated by the TOR pathway.

When looking at the time course of development, it appears that already young seedlings are affected in photosynthesis and chlorophyll accumulation. At this early time point, supplementation with uridine can rescue the phenotype (Figure 5). However, after longer periods of nucleotide limitation, phenotypes become manifest as reflected in thinner leaves and altered chloroplast ultrastructure in *ATC* mutants. In addition, a high number of plastoglobuli were observed in *ATC* lines. These electron-dense bodies within chloroplasts are involved in thylakoid lipid remodeling (Rottet et al., 2015), and chlorophyll breakdown (van Wijk and Kessler, 2017).

Photosynthetic yield was clearly reduced in both *ATC* lines, but not in *DHODH* lines (Figure 7). This reduced photosynthetic efficiency may result from photoinhibition, reflecting a light-induced damage of the PSII reaction center, followed by free radical-induced damage of other photosynthetic components (Figure 7) (Järvi et al. 2015). Indeed, we also detected an increased expression of ROS marker genes (Figure 7) and higher ROS levels, as determined by NBT and DAB staining (Figure 7). Furthermore, the ROS scavenging system was affected, as dihydro-ascorbate levels were higher in both *ATC* lines. NADPH recycling is low under conditions of reduced photosynthetic yield, but the levels of NADH and NADPH are depleted further by the reduction of oxidized ascorbate via the Foyer-Asada-Halliwell pathway (Foyer and Noctor, 2011). These combined effects may thus explain the strongly decreased levels of the NADPH and NADH pools (Figure 2).

How a smaller pool of pyrimidine nucleotides in *ATC* knockdown lines and a low energy level led to photoinhibition remains an open question. We hypothesize that a low content of pyrimidine nucleotides and the resulting imbalance in purine nucleotides will negatively affect RNA synthesis, especially that of ribosomal RNAs, since they represent the main sink for nucleotides (Busche et al., 2020). Similar effects have been observed in cytidine triphosphate synthase 2 knockdown mutants (Alamdari et al., 2021; Bellin et al., 2021). Indeed, nucleotide availability can limit ribosome biogenesis (Brunkard 2020) and a resulting low translation efficiency in the chloroplast will have major consequences on the function of the organelle, especially for proteins with a high turnover rate. The most prominent candidate here is D1 from PSII. Insufficient D1 recycling will lead to photoinhibition (Järvi et al., 2015) and ROS accumulation, as observed in *ATC* lines (Figure 7).

Inefficient biosynthesis or repair of photosynthetic proteins (such as integral membrane proteins of the electron transport chain) may drive the observed alteration in chloroplast ultrastructure seen in *ATC* lines, with less dense grana stacks and loose thylakoid structures (Figure 4, Järvi et al., 2015). Reduced photosynthetic yield and the activation of ROS detoxification systems may then lead to a low energy state. Furthermore, a low NADPH pool might not be able to redox-activate the thylakoid ATP-synthase (whose protein levels decreased in both *ATC* lines in addition (Figure 7), thus further exacerbating the low energy state and impairing plant growth (Carrillo et al., 2016).

DHODH knockdown lines exhibited reduced growth and seed production, as well as decreased CO_2_ assimilation and respiration. By contrast, these lines had close to normal chloroplast ultrastructure, maximal photosynthetic yield, and nucleotide levels (Figure 2). However, DHODH knockdown lines showed specific responses as well, such as a clear delay in germination. Respiration is the sole energy source during germination. It is therefore tempting to speculate that DHODH may play a regulatory role during respiration. However, current experimental evidence supports an opposite hierarchy, as hypoxic cells with reduced respiration are characterized by pyrimidine deficiency (Wang et al., 2019; Bajzikova et al., 2019). In Arabidopsis, seed hydration is immediately followed by oxygen consumption but also a gradual accumulation of succinate and lactate, possibly reflecting partial hypoxia in embryo tissues after the onset of germination (Nietzel et al., 2019). It is thus possible that germination may be partially characterized by low pyrimidine availability; we would expect DHODH knockdown lines to exacerbate this physiological state.

Interestingly, a connection between pyrimidine metabolism and mitochondrial function and morphology was established in human and mouse cell lines, where DHODH inhibitors induced an accumulation of mitochondrial fusion proteins and caused mitochondrial elongation (Löffler et al., 2020). In line with this observation, we detected morphological alterations in mitochondria from *DHODH* knockdown lines.

Whether DHODH is involved in mitochondrial processes other than oxidizing dihydroorotate during *de novo* pyrimidine synthesis was tested in a study with *Toxoplasma gondii*. This unicellular organism has a DHODH that belongs to the same subfamily as plant DHODHs and is also coupled to the mitochondrial respiratory chain through ubiquinone-mediated oxidation of dihydroorotate (Triana et al., 2016). Whereas supplementation of the growth medium with the salvage substrate uracil did not allow a DHODH loss of function mutant to survive, complementation with a fully catalytically inactive DHODH mutant version was possible. This result indicated that *T. gondii* DHODH is required for a second essential function (Triana et al., 2016). Complementation approaches like those tried in *T. gondii* are still out of reach in plants.

## Material and Methods

### Plant growth

For DNA isolation, tissue collection and phenotypic inspection, wild-type and transgenic *Arabidopsis thaliana* (L.) Heynh. plants (ecotype Columbia) were used throughout. Plants were grown in standardized ED73 (Einheitserde und Humuswerke Patzer) soil or on agar plates under long day conditions in a 14 h light and 10 h dark regime (120 μmol quanta m^−2^ s^−1^, temperature 22°C, humidity 60%) Illumination was done with LED light (Valoya NS1, Valoya, Finnland). For growth experiments on sterile agar plates, surface-sterilized seeds were grown on half strength MS, supplemented with 0.1% (w/v) sucrose. Prior to germination, seeds were incubated for 24 h in the dark at 4°C for imbibition (Weigel and Glazebrook, 2002). If not stated otherwise, plant material was harvested in the middle of the light period and frozen in liquid nitrogen for further use.

### Construction of DHODH knock down plants

For generation of DHODH (*pyrD;* At5g23300) RNAi lines, the procedure described in Wesley et al. (2001) was used. To incorporate the *pyrD* gene fragment in antisense orientation DHODH antisense_fwd and rev primers (Supplemental Table S1) were used. The XbaI/ BamH1 digested PCR fragment was integrated into the corresponding sites of the pHannibal vector. For amplification of the *pyrD* sense fragment DHODH_sense_fwd and rev primers (Supplemental Table S1) were used, and subsequently the PCR product was introduced into pHannibal via XhoI/ EcoRI sites. The gene expression cartridge including a CMV-35S promotor was then introduced into the NotI site of the binary vector pART27 (Gleave, 1992). All constructs used for Arabidopsis transformation by floral dip (Narusaka et al., 2010) were previously transformed into *A. tumefaciens* strain GV3101 (pMP90; (Furini et al., 1994)). Several independent, transformed lines were obtained exhibiting different DHODH transcript levels. Two of these (*dhodh#*1 and *dhodh#2*) were selected for further analysis.

### RNA extraction and gene expression analysis

Leaf material of soil grown plants was collected and homogenized in liquid nitrogen prior to extraction of RNA with the Nucleospin RNA Plant Kit (Macherey-Nagel, Düren, Germany) according to the manufacturer’s advice. RNA purity and concentration were quantified using a Nanodrop spectrophotometer. Total RNA was transcribed into cDNA using the qScript cDNA Synthesis Kit (Quantabio, USA). qPCR was performed using the quantabio SYBR green quantification kit (Quantabio) on PFX96 system (BioRad, Hercules, CA, USA) using specific primers Supplementary method S1, and At2g3760 (Actin) was used as reference gene for transcript normalization. At least three biological replicates were analyzed. Mean values and standard errors were calculated from at least three biological replicates.

### Global transcript analysis

RNA was isolated from overground tissue of six-week-old plants in three biological replicates, as described above. Cloning and sequencing was performed at Novogene (China/UK). Illumina sequencing was performed and read quality checked by Hisat2. All data shown exhibit adjusted p-values < 0.05. Differentially expressed genes were analyzed by DESeq2 ((Love *et al*., 2014).

### Protein extraction and immunoblotting

Leaf extract of wild type and mutants was prepared by homogenizing leaf material in extraction buffer (50 mM HEPES-KOH, pH 7.2, 5 mM MgCl2, 2 mM phenylmethylsulfonyl fluoride (PMSF)) on ice. This homogenous extract was centrifuged for 10 min, 20,000g and 4°C. The supernatant was collected and stored on ice until use. For immunoblotting 15 µg of a protein extract from Arabidopsis leaves separated in a 15% SDS-PAGE gel were transferred onto a nitrocellulose membrane (Whatman, Germany) by wet blotting. The membrane was blocked in phosphate-buffered saline plus 0.1% [v/v] Tween 20 (PBS-T) with 3% milk powder for 1 h at room temperature, followed by three washes of 10 min in PBS-T. Then, the membrane was incubated with a rabbit polyclonal antiserum raised against recombinant ATC (Eurogentec, Belgium) for 1 h, followed by three washes with PBS-T. Antibodies against photosynthetic proteins (PsaA #AS06172100, PsbA #AS05084, PsbD #AS06146, lhcb2 #AS01003, lhca1 #AS01005, AtpB #AS05085, cytb6-PetB #AS148169) were purchased from Agrisera (Vännäs, Sweden) Next, the membrane was incubated for 1 h with a horseradish peroxidase (HPR) conjugated anti-rabbit antibody (Promega, Walldorf, Germany) diluted in PBS-T with 3% milk powder. The result was visualized by chemiluminescence using the ECL Prime Western blotting reagent (GE Healthcare) and a Fusion Solo S6 (Vilber-Lourmat) imager.

### Chlorophyll analysis

Photosynthetic pigments were extracted from ground leave tissue with 90% acetone/ 10% 0.2 M Tris/HCl pH 7.5 for 48h at 4°C in the dark. Chlorophyll was measured by the absorbance of the supernatant at 652 nm. The quantification was performed as described by Arnon (1949).

### Generation of constructs and staining for GUS Activity

For the histochemical localization of promoter activity of *PYRB* and *PYRD*, a 965 bp upstream fragment of *PYRB* (ATC) was inserted to pBGWFS7 (Karimi et al., 2002) using the primers ATC_gus_fwd and ATC_gus_rev and a 1140 bp upstream fragment of *PYRD* (DHODH) was inserted to pGPTV (Becker et al., 1992) using the primers DHODH_gus_fwd and DHODH_gus_rev (Supplemental Table 1). The resulting constructs were transformed in *Agrobacterium* strain GV3101. Transformation of *Arabidopsis* was conducted according to the floral dip method (Clough and Bent, 1998). carrying transcriptional fusions of the GUS open reading frame with promoters of both genes. For each construct 5 independent primary transformed (F2) lines were inspected. Tissue from transgenic plants was collected in glass vials, filled with ice-cold 90% acetone, and incubated for 20 min at room temperature. Subsequently, the samples were stained according to standard protocols (Weigel and Glazebrook, 2002).

### Germination assays and root growth tests

Seed germination was analyzed with three petri dishes per genotype (each with 40 seeds) and 3 replications of the complete experiment. Seeds were grown on agar plates starting at the onset of light. After indicated time points seeds were inspected for radicle protrusion. For root growth seeds were treated as indicated above and grown vertically on square (120 × 120 mm) petri plates. 20 seeds per genotype were inspected in parallel and the experiment was repeated 3 times. Root length of seven days old seedlings was measured after scanning of agar plates with help of ImageJ software.

### Light- and electron microscopy

For image analysis of freshly prepared siliques, a Keyence VHX-5000 digital microscope (Keyence Germany GmbH, Neu-Isenburg, Germany) has been used. For histological and ultrastructural examinations, combined conventional and microwave-assisted fixation, substitution, and resin embedding of 2mm^2^ leaf cuttings were performed using a PELCO e BioWave® Pro+ (TedPella, Redding, CA, USA), according to Supplementary method S2. Therefore 4-6 cuttings of the central part of 2 different rosette leaves of at least 4 different WT and mutant plants were used. Sectioning of resin blocks, histological staining, light- and electron microscopical analysis has been carried out as described previously (Daghma et al., 2011).

### PAM measurements

A MINI-IMAGING-PAM fluorometer (Walz Instruments, Effeltrich, Germany) was used for *in vivo* chlorophyll A light curve assays on intact, 6-week-old dark-adapted plants using standard settings (Schreiber et al., 2007). Measurements were performed with eight plants per line in light curves recorded by incrementally increasing light pulses with intensity from PAR (µmol photons m^-2^ s^-1^) 0 to PAR 726 in 14 steps.

### Gas exchange measurements

Plants were grown for 6 weeks on soil in a 10h/ 14h light and dark regime. Gas exchange-related parameters were analyzed with a GFS-3000 system (Heinz Walz, Effeltrich, Germany). Measurements were performed with six plants per condition and each plant was measured three times (technical replicates). Individual plants were placed in a whole plant gas exchange cuvette and CO_2_-assimilation rate, respiration, leaf CO_2_ concentration, and stomatal conductance were recorded. Temperature, humidity, and CO_2_ concentrations of the cuvette were set to the condition’s plants were grown at. Light respiration was measured at PAR 125 and dark respiration at PAR 0 over a time of 1 min for each plant. Each plant was measured three times with 30 seconds intervals between measurement to allow leaves to return to the stabilized value.

### Superoxide and H_2_O_2_ staining

Superoxide and H_2_O_2_ staining were visually detected with nitro blue tetrazolium (NBT) and 3,3’-diaminobenzidine (DAB). In situ detection of O^2−^ was performed by treating plants with NBT as previously described by Wohlgemuth et al. (2002). *A. thaliana* leaves were vacuum-infiltrated with 0.1% NBT 50 mM potassium phosphate buffer (pH 7.8) and 10 mM sodiumazide for 20 minutes and incubated for 1 h at room temperature. Stained leaves were boiled in 95% Ethanol for 15 minutes and photographed. Detection of H_2_O_2_ was performed by treating plants with DAB-HCl as previously described by Fryer et al. (2002). Leaves were vacuum-infiltrated with 5 mM DAB-HCl, pH 3, for 20 min, and incubated in the same solution for at least 8 hours overnight. Stained leaves were boiled in an ethanol:acetic acid:glycerol (3:1:1) solution under the hood until they turned transparent and were later photographed.

### Metabolite Extraction and Quantification LC-MS

For the metabolite profiling, the freeze-dried and homogenized samples were extracted according to Schwender et al., (2015). Untargeted profiling of anionic central metabolites was performed using the Dionex-ICS-5000+HPIC the ion chromatography system (Thermo Scientific) coupled to a Q-Exactive Plus hybrid quadrupol-orbitrap mass spectrometer (Thermo Scientific). The detailed chromatographic and MS conditions are described in the Supplementary Method S3. The randomized samples were analyzed in full MS mode. The data-dependent MS-MS analysis for the compound identification was performed in the pooled probe, which also was used as a quality control (QC).

The batch data was processed using the untargeted metabolomics workflow of the Compound Discoverer 3.0 software (Thermo Scientific). The compounds were identified using the inhouse library, as well as a public spectral database mzCloud and the public databases KEGG, NIST and ChEBI via the mass- or formula-based search algorithm. The P-values of the group ratio were calculated by ANOVA and a Tukey-HCD post hoc analysis. Adjusted P-values were calculated using Benjamini-Hochberg correction. Untargeted profiling of amino acids and other cationic metabolites was performed using the Vanquish Focused ultra-high-pressure liquid chromatography (UHPLC) system (Thermo Scientific) coupled to a QExactive Plus mass spectrometer (Thermo Scientific). The detailed chromatographic and MS conditions are described in the Supplementary Method S2. The batch processing and compound identification workflow was essentially the same as described for the IC-MS-based untargeted profiling.

## ACCESSION NUMBERS

ATC (*pyrB;* At3g20330); DHODH (*pyrD;* At5g23300)

## ACKNOWLEDGEMENT

We thank Marion Benecke, Claudia Riemey and Kirsten Hoffie (IPK Gatersleben) for technical assistance with sample preparation for histology and electron microscopy. We thank Hardy Rolletscheck for advice and support in metabolite analysis. Furthermore, we are indebted to Monika Löffler and Wolfgang Knecht for fruitful discussions and critical reading of the manuscript. This work was funded by DFG grants (CRC Transregio TRR175, A03 to J.M. and B08 to T.M).

## SUPPLEMENTAL DATA

Supplemental Table S1. Primers used in this study

Supplemental Table S2. Protocol for preparation of leave cuttings for histological and ultrastructural analysis

Supplemental Table S3. Chromatographic and mass spectrometry conditions for the untargeted metabolite analysis

Supplemental Figure S1. Phenotype of ATC and DHODH mutant plants at flowering time Supplemental Figure S2. Heatmap of relative changes in quantities of selected metabolites

Supplemental Figure S3. Lists of DEGs sorted to selected pathways

Supplemental Figure S4. Histochemical staining showing of *ATC::GUS* and *DHODH::GUS* lines

Supplemental Figure S5. Histochemical staining showing of *ATC::GUS* and *DHODH::GUS* during leaf maturation

Supplemental Figure S6. Analysis of leaf ultrastructure: leaf thickness and organelle area

## LITERATURE CITED

Alamdari K, Fisher KE, Tano DW, Rai S, Palos K, Nelson ADL., et al. (2021). Chloroplast quality control pathways are dependent on plastid DNA synthesis and nucleotides provided by cytidine triphosphate synthase two. New Phytol. doi: 10.1111/nph.17467

Arnon DI (1949). Copper enzymes in isolated chloroplasts. Polyphenoloxidase in Beta vulgaris. Plant Physiol. 24: 1–15

Bajzikova M., Kovarova J, Coelho AR, Boukalova S, Oh S, Rohlenova K, Svec D, et al. (2019). Reactivation of dihydroorotate dehydrogenase-driven pyrimidine biosynthesis restores tumor growth of respiration-deficient cancer cells. Cell metabol. 29: 399–416

Becker D, Kemper E, Schell J, Masterson R (1992). New plant binary vectors with selectable markers located proximal to the left T-DNA border. Plant Mol. Biol. 20: 1195–1197

Bellin L, Caño-Ochoa D, Velázquez-Campoy A, Möhlmann T, Ramón-Maiques S (2021). Mechanisms of feedback inhibition and sequential firing of active sites in plant aspartate transcarbamoylase. Nat. commun. 12: 1–13

Bellin L, Scherer V, Dörfer E, Lau A, Vicente AM, Meurer J, Hickl D, Möhlmann T (2021). Cytosolic CTP production limits the establishment of photosynthesis in Arabidopsis. Front. Plant Sci. 12: 789189 doi: 10.3389/fpls.2021.789189

Brunkard J O (2020). Exaptive Evolution of Target of Rapamycin Signaling in Multicellular Eukaryotes. Devel. Cell

Brychkova G, Alikulov Z, Fluhr R, Sagi M (2008). A critical role for ureides in dark and senescence-induced purine remobilization is unmasked in the Atxdh1 Arabidopsis mutant. Plant J. 54: 496–509

Busche M, Scarpin MR, Hnasko R, Brunkard JO (2021). TOR coordinates nucleotide availability with ribosome biogenesis in plants. Plant Cell 33: 1615–1632

Carrillo LR, Froehlich JE, Cruz JA, Savage LJ, Kramer DM (2016). Multi-level regulation of the chloroplast ATP synthase: the chloroplast NADPH thioredoxin reductase C (NTRC) is required for redox modulation specifically under low irradiance. Plant J. 87: 654–663

Christopherson RI, Szabados E (1997). Nucleotide biosynthesis in mammals. PORTLAND PRESS RESEARCH MONOGRAPH, 315–335

Clough SJ, Bent AF (1998). Floral dip: a simplified method for Agrobacterium-mediated transformation of Arabidopsis thaliana. Plant J. 16: 735–743.

Chen CT, Slocum, RD (2008). Expression and functional analysis of aspartate transcarbamoylase and role of de novo pyrimidine synthesis in regulation of growth and development in Arabidopsis. Plant Physiol. Biochem. 46: 150–159

Daghma DS, Kumlehn J, Melzer M (2011). The use of cyanobacteria as filler in nitrocellulose capillaries improves ultrastructural preservation of immature barley pollen upon high pressure freezing. J. Microsc. 244: 79–84

Dahncke K, Witte CP (2013). Plant purine nucleoside catabolism employs a guanosine deaminase required for the generation of xanthosine in Arabidopsis. Plant Cell 25: 4101–4109.

Del Cano-Ochoa F, Ramon-Maiques S. (2021). Deciphering CAD: Structure and function of a mega-enzymatic pyrimidine factory in health and disease. Prot. Sci. 30: 1995–2008

Diehl FF, Miettinen TP, Elbashir R, Nabel CS, Darnell AM, Do BT, Manalis SR, Lewis CA, Vander Heiden MG (2022). Nucleotide imbalance decouples cell growth from cell proliferation. Nat. Cell Biol. 24: 1252–1264

Doremus HD, Jagendorf AT (1985). Subcellular localization of the pathway of de novo pyrimidine nucleotide biosynthesis in pea leaves. Plant Physiol. 79: 856–861

Dyson BC, Miller MA, Feil R, Rattray N, Bowsher CG, Goodacre R, Lunn JE, Johnson GN (2016). FUM2, a Cytosolic Fumarase, Is Essential for Acclimation to Low Temperature in Arabidopsis thaliana. Plant Physiol. 172: 118–127

Foyer CH, Noctor G (2011). Ascorbate and glutathione: the heart of the redox hub. Plant Physiol. 155: 2–18

Fryer MJ, Oxborough K, Mullineaux PM, Baker NR (2002). Imaging of photo-oxidative stress responses in leaves. J. Exp. Bo.t 53: 1249–1254

Furini A, Koncz C, Salamini F, Bartels D (1994). Agrobacterium-mediated transformation of the desiccation-tolerant plant Craterostigma plantagineum. Plant Cell. Rep. 14: 102–106

Garavito MF, Narváez-Ortiz HY, Zimmermann BH (2015). Pyrimidine metabolism: dynamic and versatile pathways in pathogens and cellular development. J. Gen. Genet. 42: 195–205

Garcia-Molina A, Kleine T, Schneider K, Mühlhaus T, Lehmann M., Leister D (2020). Translational components contribute to acclimation responses to high light, heat, and cold in Arabidopsis. IScience, 23: 101331

Geigenberger P, Regierer B, Nunes-Nesi A, Leisse A, Urbanczyk-Wochniak E, Springer F, van Dongen JT, Kossmann J Fernie AR (2005). Inhibition of de novo pyrimidine synthesis in growing potato tubers leads to a compensatory stimulation of the pyrimidine salvage pathway and a subsequent increase in biosynthetic performance. Plant Cell 17: 2077–2088

Giermann N, Schröder M, Ritter T, Zrenner R (2002). Molecular analysis of de novo pyrimidine synthesis in solanaceous species. Plant Mol. Biol. 50: 393–403

Gillespie KM, Ainsworth EA (2007). Measurement of reduced, oxidized and total ascorbate content in plants. Nat. Protoc. 2: 871–874

Gleave AP (1992). A versatile binary vector system with a T-DNA organisational structure conducive to efficient integration of cloned DNA into the plant genome. Plant Mol. Biol. 20: 1203–1207

Järvi S, Suorsa M, Aro EM (2015). Photosystem II repair in plant chloroplasts—regulation, assisting proteins and shared components with photosystem II biogenesis. Biochim. Biophys. Acta 1847: 900–909

Kafer C, Zhou L, Santoso D, Guirgis A, Weers B, Park S, Thornburg R (2004) Regulation of pyrimidine metabolism in plants. Front. Biosci. 9: 1611–1625

Karimi M, Inzé D, Depicker A (2002). GATEWAY™ vectors for Agrobacterium-mediated plant transformation. TIPS 7: 193–195

Karve A, Moore BD (2009). Function of Arabidopsis hexokinase-like1 as a negative regulator of plant growth. J. Exp. Bot. 60: 4137–4149

Kato Y, Miura E, Ido K, Ifuku K, Sakamoto W (2009). The variegated mutants lacking chloroplastic FtsHs are defective in D1 degradation and accumulate reactive oxygen species. Plant Physiol. 151: 1790–1801

Kim H, Kelly RE, Evans DR (1992). The structural organization of the hamster multifunctional protein CAD. Controlled proteolysis, domains, and linkers. J. Biol. Chem. 267: 7177–7184

Kirch HH, Bartels D, Wei Y, Schnable PS, Wood AJ (2004). The ALDH gene superfamily of Arabidopsis. TIPS 9: 371–377

Lambrev PH, Miloslavina Y, Jahns P, Holzwarth AR (2012). On the relationship between non-photochemical quenching and photoprotection of Photosystem II. Biochim. Biophys. Acta 1817: 760–769

Lehninger AL, Nelson DL, Cox MM (1994). Prinzipien der Biochemie, H. Tschesche, ed (Heidelberg, Berlin, Oxford: Spektrum Akademischer Verlag).

Linka N, Weber AP (2010). Intracellular metabolite transporters in plants. Mol. Plant 3: 21–53

Löffler M, Carrey EA, Knecht W (2020). The pathway to pyrimidines: The essential focus on dihydroorotate dehydrogenase, the mitochondrial enzyme coupled to the respiratory chain. Nucleosides, Nucleotides and Nucleic Acids 39: 1281–1305

Love MI, Huber W, Anders S (2014). Moderated estimation of fold change and dispersion for RNA-seq data with DESeq2. Genome Biol. 15: 550

Martinussen J, Willemoës M, Kilstrup M (2011). Nucleotide Metabolism. In: M. Moo-Young (Ed.), Comprehensive Biotechnology (2 ed., Vol. 1, pp. 91–107). Elsevier. http://www.sciencedirect.com/science/referenceworks/9780080885049#ancsec1

Müller P, Li XP, Niyogi KK (2001). Non-photochemical quenching. A response to excess light energy. Plant Physiol. 125: 1558–1566

Moffatt BA, Ashihara H (2002). Purine and pyrimidine nucleotide synthesis and metabolism. The Arabidopsis Book/American Society of Plant Biologists, 1.

Moreno-Morcillo M, Grande-Garcia A, Ruiz-Ramos A, Del Cano-Ochoa F, Boskovic J, Ramon-Maiques S (2017). Structural insight into the core of CAD, the multifunctional protein leading de novo pyrimidine biosynthesis. Structure 25: 912–923 e915

Nara T, Hshimoto T, Aoki T (2000). Evolutionary implications of the mosaic pyrimidine-biosynthetic pathway in eukaryotes. Gene 257: 209–222

Narusaka M, Shiraishi T, Iwabuchi M, Marusaka Y (2010). The floral inoculating protocol: A simplified Arabidopsis thaliana transformation method modified from floral dipping. Plant Biotech. 27: 349–351

Nasr F, Bertauche N, Dufour ME, Minet M, Lacroute F (1994). Heterospecific cloning of Arabidopsis thaliana cDNAs by direct complementation of pyrimidine auxotrophic mutants of Saccharomyces cerevisiae. I. Cloning and sequence analysis of two cDNAs catalysing the second, fifth and sixth steps of the de novo pyrimidine biosynthesis pathway. Mol. Genet. Gen. 244: 23–32

Nietzel T, Mostertz J, Ruberti C, Née G, Fuchs P, Wagner S, Moseler A, Müller-Schüssele SJ, Benamar A, Poschet G et al. (2020). Redox-mediated kick-start of mitochondrial energy metabolism drives resource-efficient seed germination. Proc. Natl. Acad. Sci. 117: 741–751

Noctor G, Foyer CH (1998). Ascorbate and glutathione: keeping active oxygen under control. Annu. Rev. Plant. Biol. 49: 249–279

Ohler L, Niopek-Witz S, Mainguet SE, Möhlmann, T (2019). Pyrimidine salvage: Physiological functions and interaction with chloroplast biogenesis. Plant Physiol. 180: 1816–1828

Pracharoenwattana I, Zhou W, Keech O, Francisco PB, Udomchalothorn T, Tschoep H, Stitt M, Gibon Y, Smith SM (2010). Arabidopsis has a cytosolic fumarase required for the massive allocation of photosynthate into fumaric acid and for rapid plant growth on high nitrogen. Plant J. 62: 785–795

Queval G, Noctor G (2007). A plate reader method for the measurement of NAD, NADP, glutathione, and ascorbate in tissue extracts: application to redox profiling during Arabidopsis rosette development. Anal. Biochem. 363: 58–69

Reichard P (1988). Interactions between deoxyribonucleotide and DNA synthesis. Annu. Rev. Biochem. 57: 349–374

Rottet S, Besagni C, Kessler F (2015) The role of plastoglobules in thylakoid lipid remodeling during plant development. Biochim. Biophys. Acta 1847: 889–899

Santoso D, Thornburg, R (1998). Uridine 5′-monophosphate synthase is transcriptionally regulated by pyrimidine levels in Nicotiana plumbaginifolia. Plant Physiol. 116: 815–821

Schmid, LM, Ohler L, Möhlmann T, Brachmann A, Muiño JM, Leister D, et al. (2019). PUMPKIN, the sole plastid UMP kinase, associates with group II introns and alters their metabolism. Plant Physiol. 179: 248–264

Schreiber U, Quayle P, Schmidt S, Escher BI, Mueller JF (2007). Methodology and evaluation of a highly sensitive algae toxicity test based on multiwell chlorophyll fluorescence imaging. Biosens. Bioelectron. 22: 2554–2563

Schröder M, Giermann N, Zrenner R (2005). Functional analysis of the pyrimidine de novo synthesis pathway in solanaceous species. Plant Physiol. 138: 1926–1938

Schwender J, Hebbelmann I, Heinzel N, Hildebrandt T, Rogers A, Naik D, Klapperstück M, Braun HP, Schreiber F, Denolf P et al. (2015). Quantitative multilevel analysis of central metabolism in developing oilseeds of oilseed rape during in vitro culture. Plant Physiol. 168: 828–848.

Stitt M, Lilley RMcC, Gerhardt R, Heldt HW (1989). Metabolite levels in specific cells and subcellular compartments of plant leaves. In: Methods in Enzymology (Academic Press) 32: 518–552

Su J, Yang L, Zhu Q, Wu H, He Y, Liu Y, et al. (2018). Active photosynthetic inhibition mediated by MPK3/MPK6 is critical to effector-triggered immunity. PLoS Biol. 16: e2004122. https://doi.org/10.1371/journal.pbio.2004122

Triana MAH, Herrera DC, Zimmermann BH, Fox BA, Bzik DJ (2016). Pyrimidine pathway-dependent and-independent functions of the Toxoplasma gondii mitochondrial dihydroorotate dehydrogenase. Infection and immunity, 84: 2974–2981

Trentmann O, Mühlhaus T, Zimmer D, Sommer F, Schroda M, Haferkamp I, Keller I, Pommerrenig B, Neuhaus HE (2020). Identification of chloroplast envelope proteins with critical importance for cold acclimation. Plant Physiol. 182: 1239–1255

Ullrich A, Knecht W, Piskur J, Löffler M (2002). Plant dihydroorotate dehydrogenase differs significantly in substrate specificity and inhibition from the animal enzymes. FEBS Lett. 529: 346–350

van Wijk KJ, Kessler F (2017). Plastoglobuli: plastid microcompartments with integrated functions in metabolism, plastid developmental transitions, and environmental adaptation. Annu. Rev. Plant Biol. 68: 253–289

Wang Y, Bai C, Ruan Y, Liu M, Chu Q, Qiu L, et al. (2019). Coordinative metabolism of glutamine carbon and nitrogen in proliferating cancer cells under hypoxia. Nat. Commun. 14: 201. doi: 10.1038/s41467-018-08033-9

Weigel D, Glazebrook J (2002). Arabidopsis. A laboratory manual (New York, NY: Cold Spring Harbor Laboratory Press)

Wesley SV, Helliwell CA, Smith NA, Wang MB, Rouse DT, Liu Q, et al. (2001). Construct design for efficient, effective and high-throughput gene silencing in plants. Plant J. 27: 581–590

Williamson CL, Slocum RD (1994). Molecular cloning and characterization of the pyrB1 and pyrB2 genes encoding aspartate transcarbamoylase in pea (Pisum sativum L.). Plant Physiol. 105: 377–384

Williamson CL, Lake MR, Slocum RD (1996). A cDNA encoding carbamoyl phosphate synthetase large subunit (carB) from Arabidopsis (Accession No. U40341)(PGR96-055). Plant Physiol. 111, 1.

Witz S, Jung B, Fürst S, Möhlmann T (2012). De novo pyrimidine nucleotide synthesis mainly occurs outside of plastids, but a previously undiscovered nucleobase importer provides substrates for the essential salvage pathway in Arabidopsis. Plant Cell 24: 1549–1559

Witte CP, Herde M (2020). Nucleotide metabolism in plants. Plant Physiol. 182: 63–78

Wohlgemuth H, Mittelstrass K, Kschieschan S, Bender J, Weigel HJ, Overmyer K, et al. (2002). Activation of an oxidative burst is a general feature of sensitive plants exposed to the air pollutant ozone. Plant Cell Environ. 25: 717–726

Xiong Y, McCormack M, Li L, Hall Q, Xiang C, Sheen J (2013). Glucose–TOR signalling reprograms the transcriptome and activates meristems. Nature 496: 181–186

Zrenner R, Stitt M, Sonnewald U, Boldt R. (2006). Pyrimidine and purine biosynthesis and degradation in plants. Annu. Rev. Plant Biol. 57: 805–836

