## Supplemental data for "Nucleotide limitation results in impaired photosynthesis, reduced growth and seed yield together with massively altered gene expression"

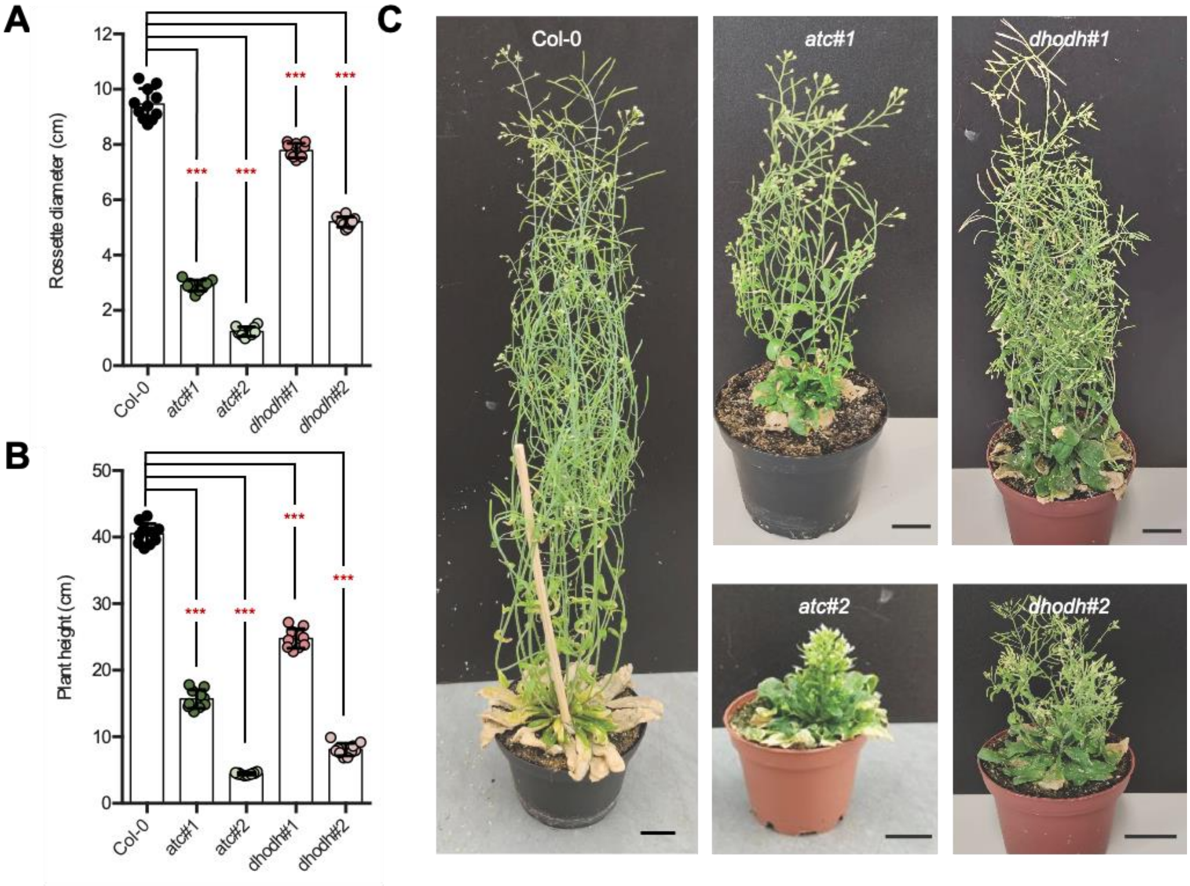


**Supplemental Figure S1 Phenotype of *ATC* and *DHODH* mutant plants at flowering time.** (**A**) Rosette diameter of 8-week-old Arabidopsis plants (n = 10). (**B**) Analysis of the maximum height of adult Col-0, *ATC* and *DHODH* mutants (n = 10). (**C**) Typical images of adult Col-0, *ATC* and *DHODH* mutants. Data points are means ± SE. Asterisks depict significant changes between the different lines referring to the WT according to one-way ANOVA followed by the Dunnett's multiple comparison test (* = $p < 0.05$, ** = $p < 0.001$, *** = $p < 0.001$).

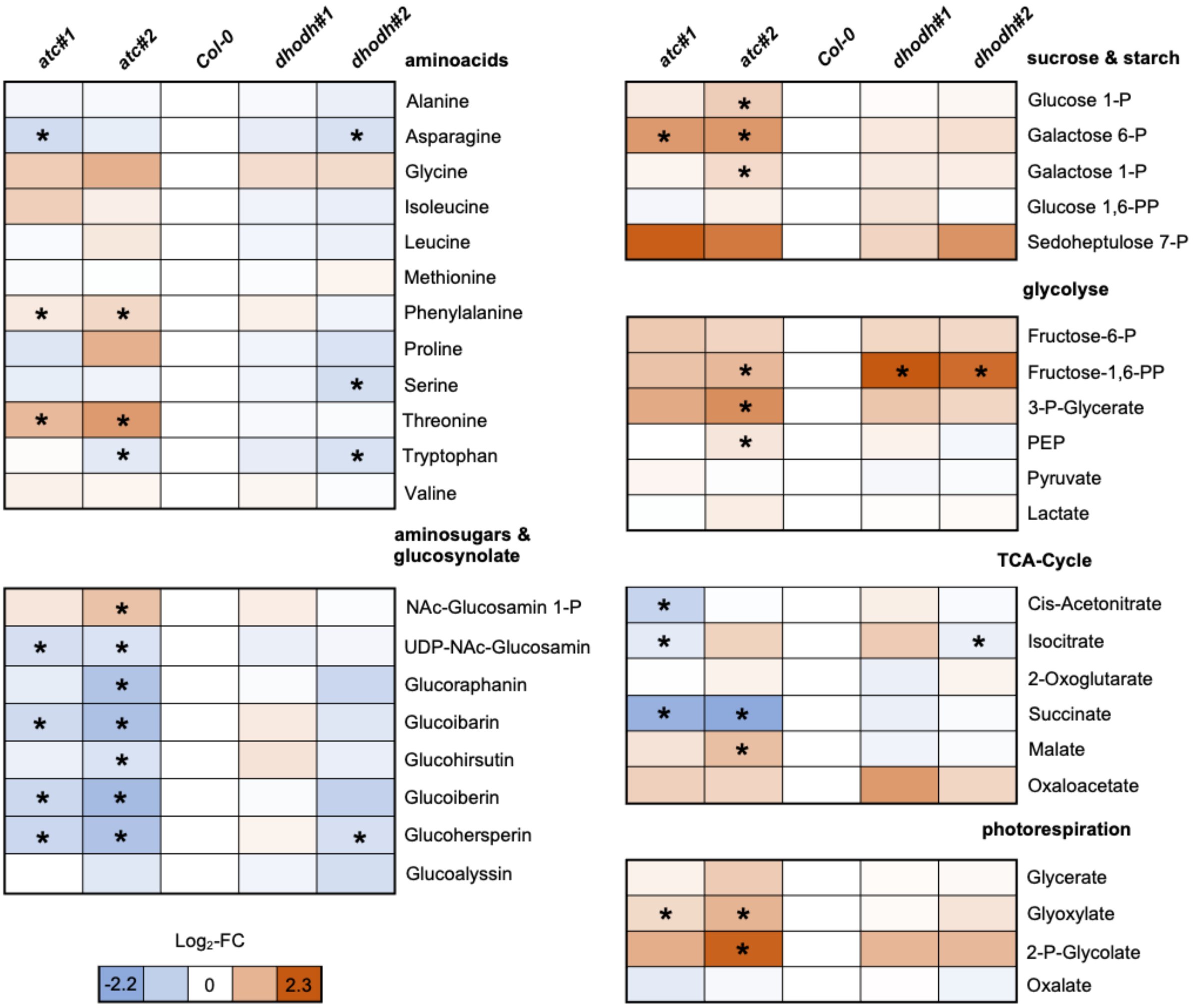


**Supplemental Figure S2. Heatmap of relative changes in quantities of selected metabolites.** $Log_2$-Foldchanges ($Log_2$-FC) are indicated by color-code. Asterisks indicate significant difference to Col-0. For determination of statistical significance (p-value < 0.05) Wilcoxon Mann-Whitney U-test was performed

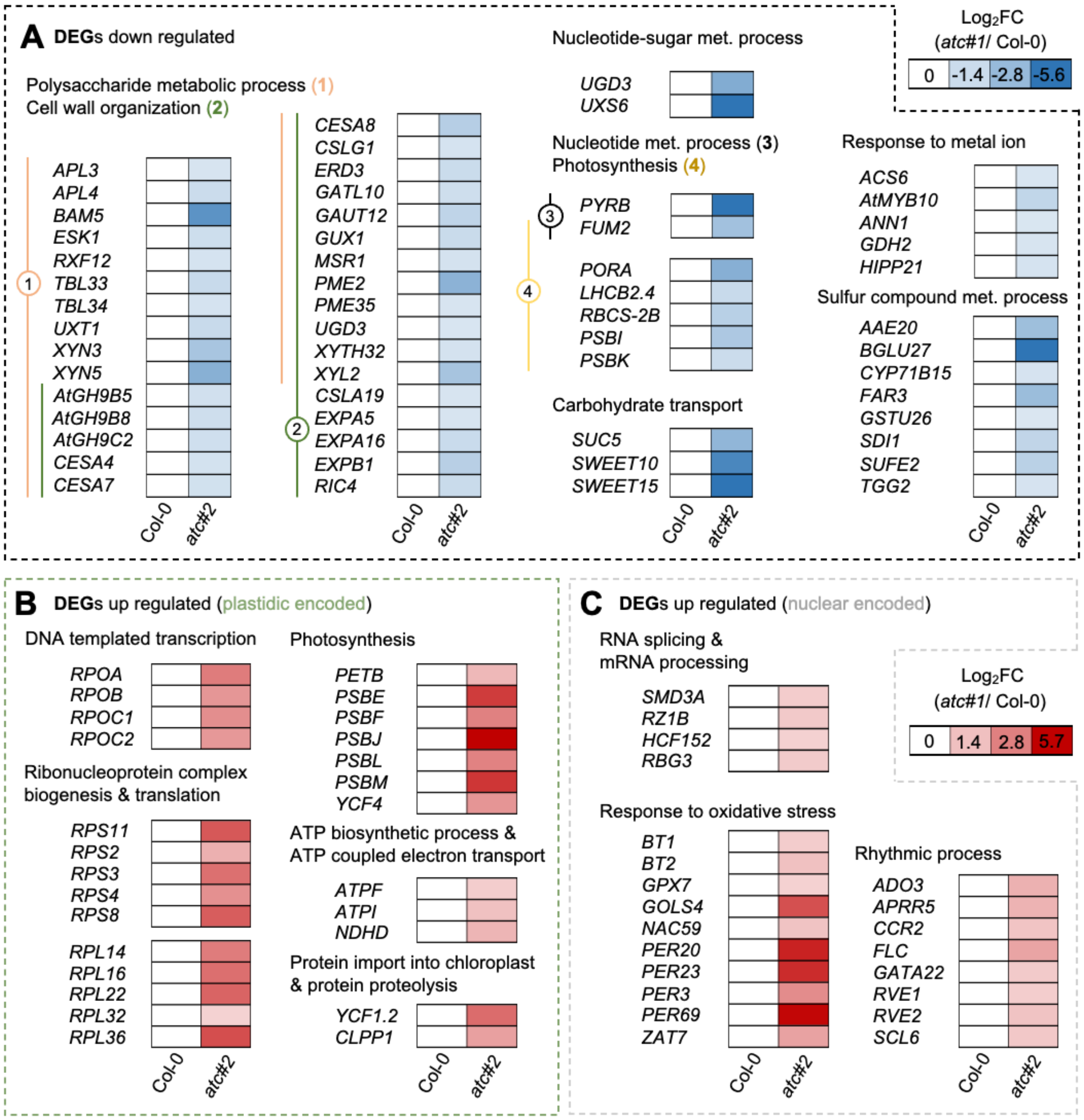


**Supplemental Figure S3. Lists of DEGs sorted to selected pathways.** Shown are DEGs in selected pathways with significant ($p_{adj.} < 0.05$) altered expression. (**A**) $Log_2$Foldchange < -1 or a (**B, C**) $Log_2$Foldchange > +1 in atc#1 knockdown mutants compared to Col-0.

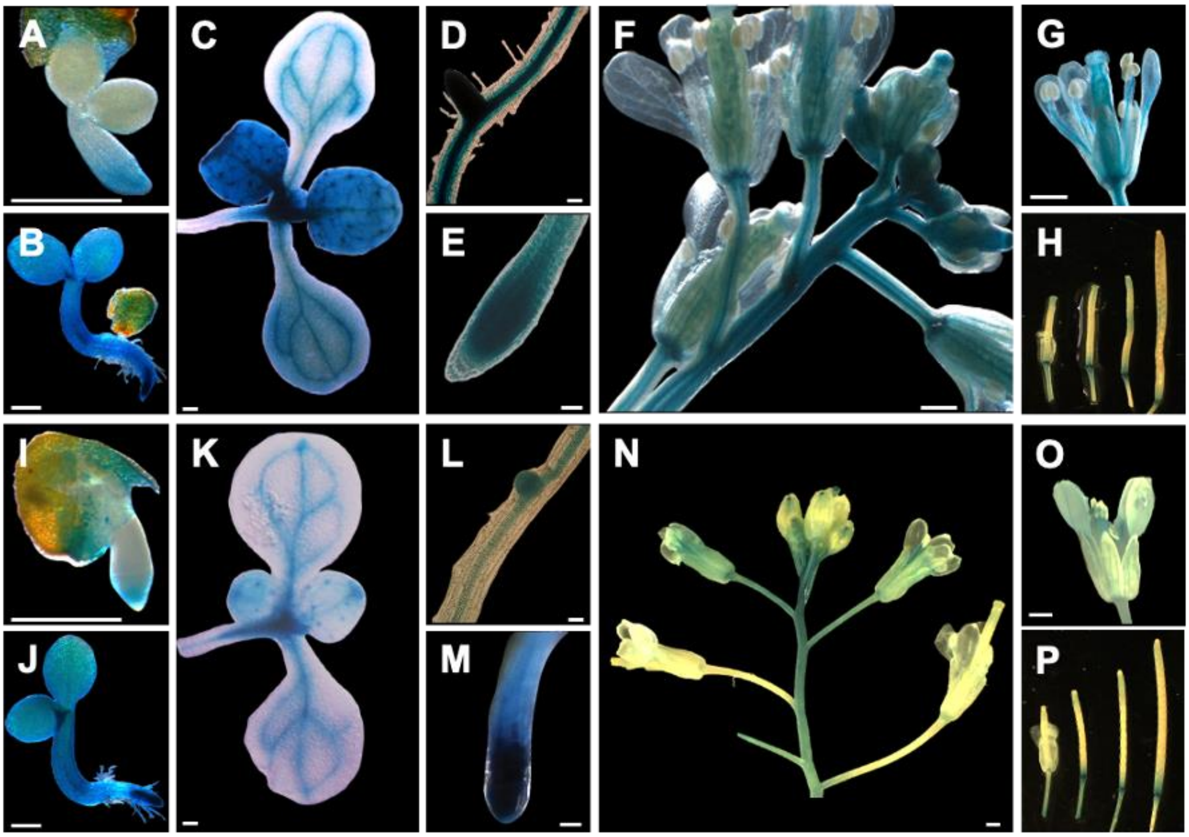


**Supplemental Figure S4. Histochemical staining of *ATC::GUS* (A-H) and *DHODH::GUS* (I-P) lines.** Images are shown for two (**A, I**) and four (**B, J**) days old seedlings. Shoots of two weeks old plants (**C, K**) corresponding lateral roots (**D, L**) and root tips (**E, M**). The inflorescence of adult plants (F, N) and in closer view flower (**G, O**) and siliques (H, P) are shown. Scale bars = 0.5 mm (A-C, F-K, N-P) or = 50 µm (D, E, L, M).

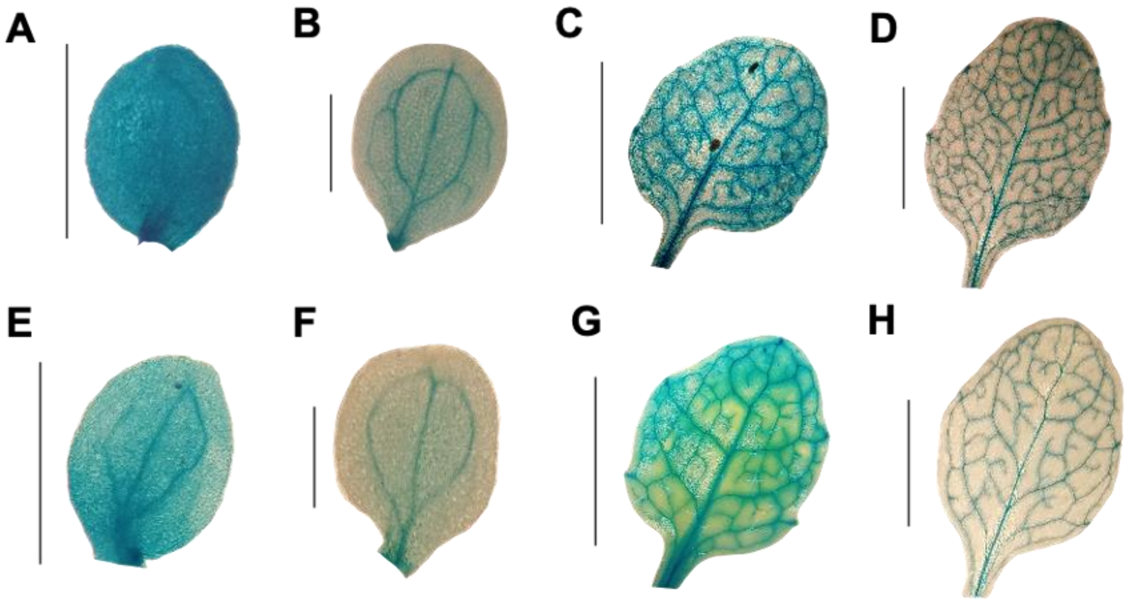


**Supplemental Figure S5. Histochemical staining of *ATC::GUS* and *DHODH::GUS* during leaf maturation.** (**A-D**) *ATC-GUS* expression (**E-H**) *DHODH-GUS* expression. (**A, E**) Cotyledones from two day old seedlings, (**B, F**) Cotyledones of four day old seedlings (**C, G**) young leaves of 15 day old plants (**D, H**) older leaves from 15 day old plant. Scale bar = 1 mm (**A, B, E, F**) = 10 mm (**C, D, G, H**).

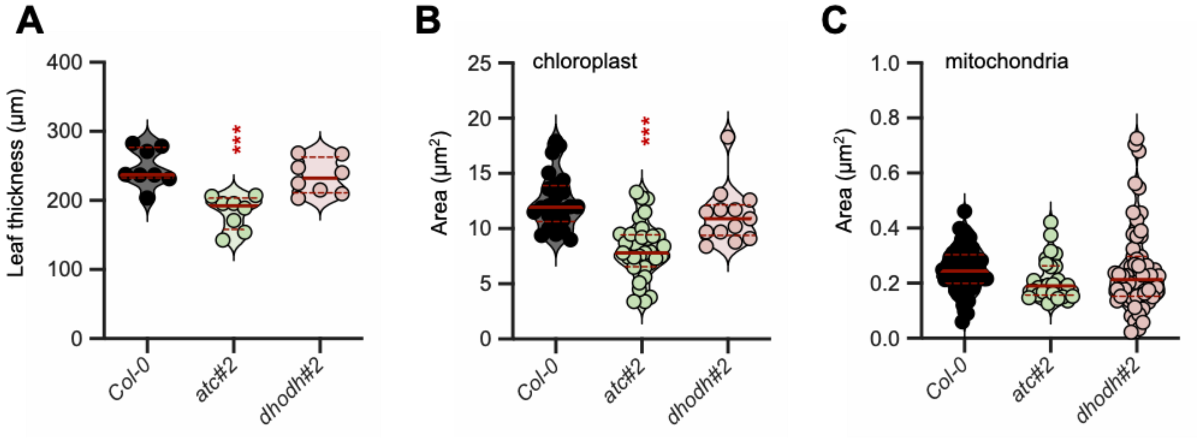


**Supplemental Figure S6. Analysis of leaf ultrastructure: leaf thickness and organelle area.** (**A**) Leaf thickness and organelle area of (**B**) chloroplast and (**C**) mitochondria in Col-0, *atc#2* and *dhodh#2* mutants. Shown are the means of biological replicates +/- standard deviation. Asterisks depict significant changes between the different lines referring to the WT according to one-way ANOVA followed by the Dunnett's multiple comparison test (*** = $p<0.001$).
